# Nap1 and Kap114 co-chaperone H2A-H2B and facilitate targeted histone release in the nucleus

**DOI:** 10.1101/2023.05.09.539987

**Authors:** Ho Yee Joyce Fung, Jenny Jiou, Ashley B. Neisman, Natalia E. Bernardes, Yuh Min Chook

## Abstract

Core histones are synthesized and processed in the cytoplasm before transport into the nucleus for assembly into nucleosomes; however, they must also be chaperoned as free histones are toxic. The importin Kap114 binds and transports histone H2A-H2B into the yeast nucleus, where Ran^GTP^ facilitates H2A-H2B release. Kap114 and H2A-H2B also bind the Nap1 histone chaperone, which is found in both the cytoplasm and the nucleus, but how Nap1 and Kap114 cooperate in H2A-H2B processing and nucleosome assembly has been unclear. To understand these mechanisms, we used biochemical and structural analyses to reveal how Nap1, Kap114, H2A-H2B and Ran^GTP^ interact. We show that Kap114, H2A-H2B and a Nap1 dimer (Nap1_2_) assemble into a 1:1:1 ternary complex. Cryogenic electron microscopy revealed two distinct Kap114/Nap1_2_/H2A-H2B structures: one of H2A-H2B sandwiched between Nap1_2_ and Kap114, and another in which Nap1_2_ bound to the Kap114•H2A-H2B complex without contacting H2A-H2B. Another Nap1_2_•H2A-H2B•Kap114•Ran^GTP^ structure reveals the nuclear complex. Mutagenesis revealed shared critical interfaces in all three structures. Consistent with structural findings, DNA competition experiments demonstrated that Kap114 and Nap1_2_ together chaperone H2A-H2B better than either protein alone. When Ran^GTP^ is present, Kap114’s chaperoning activity diminishes. However, the presence of Nap1_2_ within the Nap1_2_•H2A-H2B•Kap114•Ran^GTP^ quaternary complex restores its ability to chaperone H2A-H2B. This complex effectively deposits H2A-H2B into nucleosomes. Together, these findings suggest that Kap114 and Nap1_2_ provide a sheltered path from cytoplasm to nucleus, facilitating the transfer of H2A-H2B from Kap114 to Nap1_2_, ultimately directing its specific deposition into nucleosomes.

**Significance Statement:** Free histones are toxic and must be sequestered by other macromolecules in the cell. Nuclear import receptor Kap114 imports H2A-H2B into the nucleus while also chaperoning it. The histone chaperone Nap1 also chaperones H2A-H2B, but it is unclear how Nap1 and Kap114 cooperate to process H2A-H2B. We present biochemical and structural results that explain how Kap114, Nap1 and H2A-H2B assemble in the absence and presence of Ran^GTP^, how Nap1 and Kap114 co-chaperone H2A-H2B, and how Ran^GTP^ and Nap1 coordinate the transfer of H2A-H2B from Kap114 to assembling nucleosomes in the nucleus.

## Introduction

Formation of new nucleosomes to pack newly replicated DNA during the S-phase of the cell cycle begins with rapid synthesis of core histones H3, H4, H2A and H2B followed by their swift transport into the nucleus. Exposure of these very basic polypeptides to the cellular environment is deeply deleterious as they form toxic aggregates easily (1, 2). Therefore, core histones are thought to be always chaperoned and never free (3, 4). As H3 and H4 emerge from translating ribosomes, they are folded into H3-H4 heterodimers by heat shock proteins and folding chaperones, acetylated at several lysine side chains (5–7) and then passed to histone chaperone ASF1 and *H. sapiens* (*Hs*) Importin-4 or its *S. cerevisiae* (*Sc*) homolog Kap123 for transport across the nuclear pore complex (NPC) into the nucleus (8–11). Unfortunately, the steps of H2A-H2B biosynthesis and processing in the cytoplasm have not been delineated. However, selective ribosome profiling studies showed no association of *Sc* importins with the nascent chains of H2A and H2B (12), suggesting that H2A-H2B may follow a processing path like H3-H4’s that involves heat shock protein mediated folding and association with particular histone chaperones prior to importin-binding for nuclear import.

The heat shock proteins/folding chaperones for H2A and H2B have not been identified, but it is well-established that the folded H2A-H2B heterodimer binds the Nucleosome Assembly Protein 1 (Nap1) histone chaperone in the *Sc* cytoplasm (13–15). In human cells, H2A-H2B also binds several *Hs* Nap1 homologs (16, 17). The importin primarily responsible for H2A-H2B nuclear import is also known - *Sc* Karyopherin-114 (Kap114) or the homologous *Hs* Importin-9 (IMP9/IPO9).

*Sc* Nap1 is mostly localized to the yeast cytoplasm where it chaperones newly synthesized and folded H2A-H2B (13, 18). It also has well-known roles in the nucleus, including nucleosome assembly/remodeling, transcription, DNA repair and mitosis (14, 19–33). Genetic interaction between Nap1 and Kap114 as well as interactions with Ran binding proteins Yrb1 and Yrb2 have been reported (34). *Sc* Nap1 was also reported to be a cofactor for H2A-H2B nuclear import by Kap114 (13, 35). Both immunoprecipitation (IP) and size exclusion chromatography (SEC) of yeast cytosolic extract showed Kap114 association with both Nap1 and H2A-H2B while IP from nuclear extract showed persisting association of the three proteins in the Ran^GTP^-rich nucleus (13, 36). Therefore, Kap114, H2A-H2B and Nap1 proteins likely form a complex in the cytoplasm and remain associated in the nucleus prior to unloading the histones onto assembling nucleosomes (36). Nap1’s functions in both compartments, its interaction with Kap114 and its mostly cytoplasmic localization suggest that it may shuttle between the nucleus and the cytoplasm. According to the Saccharomyces Genome Database (SGD), Nap1, which often forms dimers or Nap1_2_, is three-fold more abundant than Kap114 (37). However, there is a lack of information about its temporal-spatial distribution in the cell, including how much and when it remains in the cytoplasm, enters the nucleus, or subsequently returns to the cytoplasm.

H2A and H2B contain disordered N- and C-terminal tails and central alpha helices that fold together into the globular H2A-H2B histone-fold domain (Fig. 1A) (38). The H2A-H2B domain has very basic surfaces that bind DNA in the nucleosome. Outside of the nucleosome, these basic surfaces participate in non-specific/aberrant interactions that produce toxic aggregates unless they are shielded (2). Crystal structures show Nap1_2_ binding asymmetrically to one H2A-H2B domain to shield its DNA-binding surfaces, explaining its function as a histone chaperone (22, 39).

**Figure 1.**
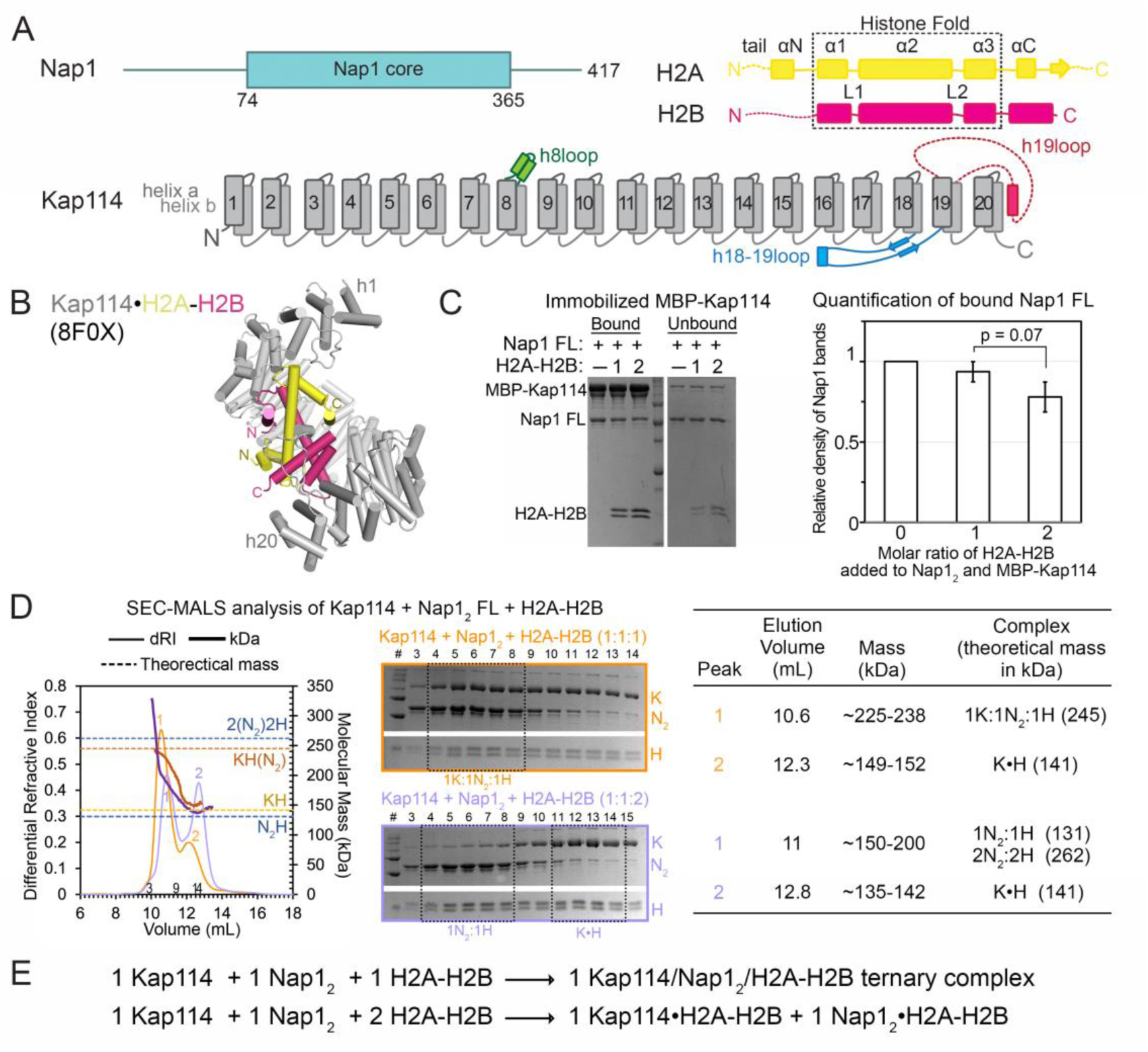
Equimolar Kap114, Nap1_2_ and H2A-H2B form a 1:1:1 complex but excess histone disrupts the ternary complex. **(A)** Organization schematics of the Nap1 (cyan), H2A (yellow), H2B (magenta) and Kap114 (gray, long loops labeled) polypeptides. **(B)** The Kap114•H2A-H2B binary complex structure (8F0X). **(C)** Pull-down assay with immobilized MBP-Kap114 (1 µM) and equimolar Nap1_2_, ± 1 or 2 molar ratio H2A-H2B; Coomassie-stained SDS-PAGE with quantified Nap1_2_ bands on the right. Controls in SI Appendix, Fig. S2. **(D)** SEC-MALS analysis of 1:1:1 (orange) or 1:1:2 (lilac) molar ratio Kap114, Nap1_2_ and H2A-H2B mixtures. Left, differential refractive index (dRI, left y-axis, thin lines) and molecular mass (kDa, right y-axis, thick lines) traces and theoretical masses of Kap114•H2A-H2B (KH), Nap1_2_•H2A-H2B (N_2_H and 2(N_2_)2H) and Kap114/Nap1_2_/H2A-H2B (K/N_2_/H) complexes marked with dashed lines. Middle panel, Coomassie-stained SDS-PAGE of peak fractions. Right, tabulated SEC-MALS results. At 1:1:1 molar ratio, the three proteins assemble into a 1 Kap114/1 Nap1_2_/1 H2A-H2B complex. No ternary complex is observed when H2A-H2B is in excess; only Kap114•H2A-H2B, Nap1_2_•H2A-H2B and 2Nap1_2_•2H2A-H2B binary complexes formed. **(E)** A summary schematic of the results in **(D)**.

The superhelical Kap114 wraps around the H2A-H2B histone-fold domain (Fig. 1B), occluding the histone’s DNA-binding surfaces and thus also acting as a histone chaperone (40, 41). The Kap114•H2A-H2B complex is unusual in its interactions with the GTPase Ran^GTP^. Most importin•cargo complexes are dissociated by Ran^GTP^ but Kap114•H2A-H2B binds Ran^GTP^ to form a stable Ran^GTP^•Kap114•H2A-H2B complex, which alters interactions between Kap114 and H2A-H2B, and facilitates histone transfer to other nuclear chaperones or to the assembling nucleosome (41).

While we understand how Nap1_2_ binds H2A-H2B and how Kap114 binds H2A-H2B, we do not know how the three proteins interact for Nap1_2_ to act as a co-factor in H2A-H2B import. We also do not know how Ran^GTP^ interacts with Kap114, Nap1_2_ and H2A-H2B in the nucleus or how their interactions affect nucleosome assembly. Here, to address these questions, we used biochemical analyses and cryo-EM structure determination to understand how Kap114, Nap1_2_ and H2A-H2B might assemble in the cytoplasm and nucleus. We showed by cryogenic electron microscopy (cryo-EM) their assembly into a 1:1:1 Kap114/Nap1_2_/H2A-H2B ternary complex that adopts two different arrangements, both showing extensive sequestration and occlusion of H2A-H2B DNA-binding interfaces by Kap114 and Nap1_2_. The structural data is consistent with co-chaperoning indicated by DNA competition assays. Analysis in the presence of Ran^GTP^ to mimic conditions in the nucleus showed the GTPase binding to form a quaternary Nap1_2_•H2A-H2B•Kap114•Ran^GTP^ complex where H2A-H2B remains sequestered by Kap114 and Nap1_2_. DNA competition and nucleosome assembly assays confirmed that in the presence of Ran^GTP^, Kap114 and Nap1_2_ continue to shield H2A-H2B from non-specific aggregation with DNA while they cooperate to transfer H2A-H2B effectively and specifically into nucleosomes.

## Results and Discussion

### Kap114 binds simultaneously to a H2A-H2B heterodimer and a Nap1 dimer

Biochemical and structural studies of binary Kap114-H2A-H2B and Nap1_2_-H2A-H2B interactions, and pull-down studies of Kap114 and Nap1 from yeast lysates have been reported (13, 22, 36, 41). First, we confirmed the previous findings using analytical ultracentrifugation (AUC), size exclusion chromatography multi-angle light scattering (SEC-MALS) and pull-down binding assays with recombinant Kap114, Nap1 and H2A-H2B (SI Appendix, Fig. S1). We showed that Nap1 alone is a mixture of dimers and tetramers (SI Appendix, Fig. S1B), and Nap1 binds H2A-H2B to form a mixture of Nap1 dimer and tetramers bound to one to two copies of H2A-H2B (SI Appendix, Fig. S1C). Nap1 binds Kap114 to form a 1:1 Kap114:Nap1_2_ complex (SI Appendix, Fig. S1A and B).

Next, we focused on interactions between the three proteins. Immobilized MBP-Kap114 pulled down Nap1 in the absence/presence of H2A-H2B; Kap114 also pulled down H2A-H2B in the absence/presence of Nap1 (Fig. 1C and SI Appendix, Fig. S2A-C). SEC-MALS analysis showed the three proteins forming a 1:1:1 Kap114/Nap1_2_/H2A-H2B complex (Fig. 1D, and SI Appendix, Fig. S2D). Interestingly, molar excess of H2A-H2B slightly decreased the amount of Nap1 pulled down by MBP-Kap114 beads (Fig. 1C and SI Appendix, Fig. S2B). SEC-MALS analysis of 1:1 Kap114:Nap1_2_ in the presence of two-fold excess H2A-H2B showed two peaks that match the binary Nap1•H2A-H2B and Kap114•H2A-H2B complexes (Fig. 1D, and SI Appendix, Fig. S1C), consistent with destabilization of the ternary complex seen in the pull-down experiment.

In summary, equimolar Kap114, Nap1_2_ and H2A-H2B assemble into a stable 1:1:1 ternary complex (Fig. 1E). Excess H2A-H2B destabilizes the ternary complex, dissociating it into binary Kap114•H2A-H2B and Nap1_2_•H2A-H2B complexes (Fig. 1E). Ternary Kap114/Nap1_2_/H2A-H2B interactions, such as in the cytoplasm, may be most stable when all H2A-H2B heterodimers are adequately chaperoned.

### Structure determination of the Kap114/Nap1_2_/H2A:H2B ternary complex

Because the Nap1_2_•H2A-H2B complex likely forms before encountering Kap114 (5, 11), we prepared a cryo-EM sample by first assembling Nap1_2_ core•H2A-H2B with excess histones to ensure binding, then adding Kap114, immediately followed by mild crosslinking (SI Appendix, Fig. S3A). After SEC, the sample remained heterogenous, a characteristic evident in the cryo-EM dataset that produced ∼1 million particles: 38.7% are Nap1_2_, 36.0% Kap114•H2A-H2B, and 25.3% ternary complex of Kap114, Nap1_2_ and H2A-H2B (SI Appendix, Fig. S3B). The smaller population of ternary complex is likely due to destabilization by excess H2A-H2B.

The Nap1_2_ and Kap114•H2A-H2B particles produced cryo-EM maps/structures that are very similar to the respective crystal and cryo-EM structures in the Protein Data Bank or PDB (SI Appendix, Fig. S4 and Table S1) (41). Particles of the ternary Kap114/Nap1_2_/H2A-H2B complex were classified into two evenly divided classes that produced two high-resolution maps and structures (Fig. 2A-F and SI Appendix, Fig. S5 and Table S1). The first is a 3.2 Å resolution map of a Nap1_2_ bound to a Kap114•H2A-H2B complex where Nap1_2_ contacts the importin but not the histone; we termed this structure Nap1_2_•Kap114•H2A-H2B (Fig. 2A and B). The second is a 3.5 Å resolution map of Kap114 and Nap1_2_ interacting with the same H2A-H2B; we termed the structure Nap1_2_•H2A-H2B•Kap114 (Fig. 2C and D). Local resolution of the Nap1_2_ region in both maps is poor, at ∼ 6-9 Å, suggesting dynamic Nap1_2_ in both structures (SI Appendix, Fig. S5A and E). Nevertheless, local refinement improved the focused maps at the Nap1_2_ regions, allowing us to dock in the histone chaperone. We emphasize the distinct protein arrangements in the two structures by highlighting ‘Kap114’ in bold: Nap1_2_•**Kap114**•H2A-H2B versus Nap1_2_•H2A-H2B•**Kap114**.

**Figure 2.**
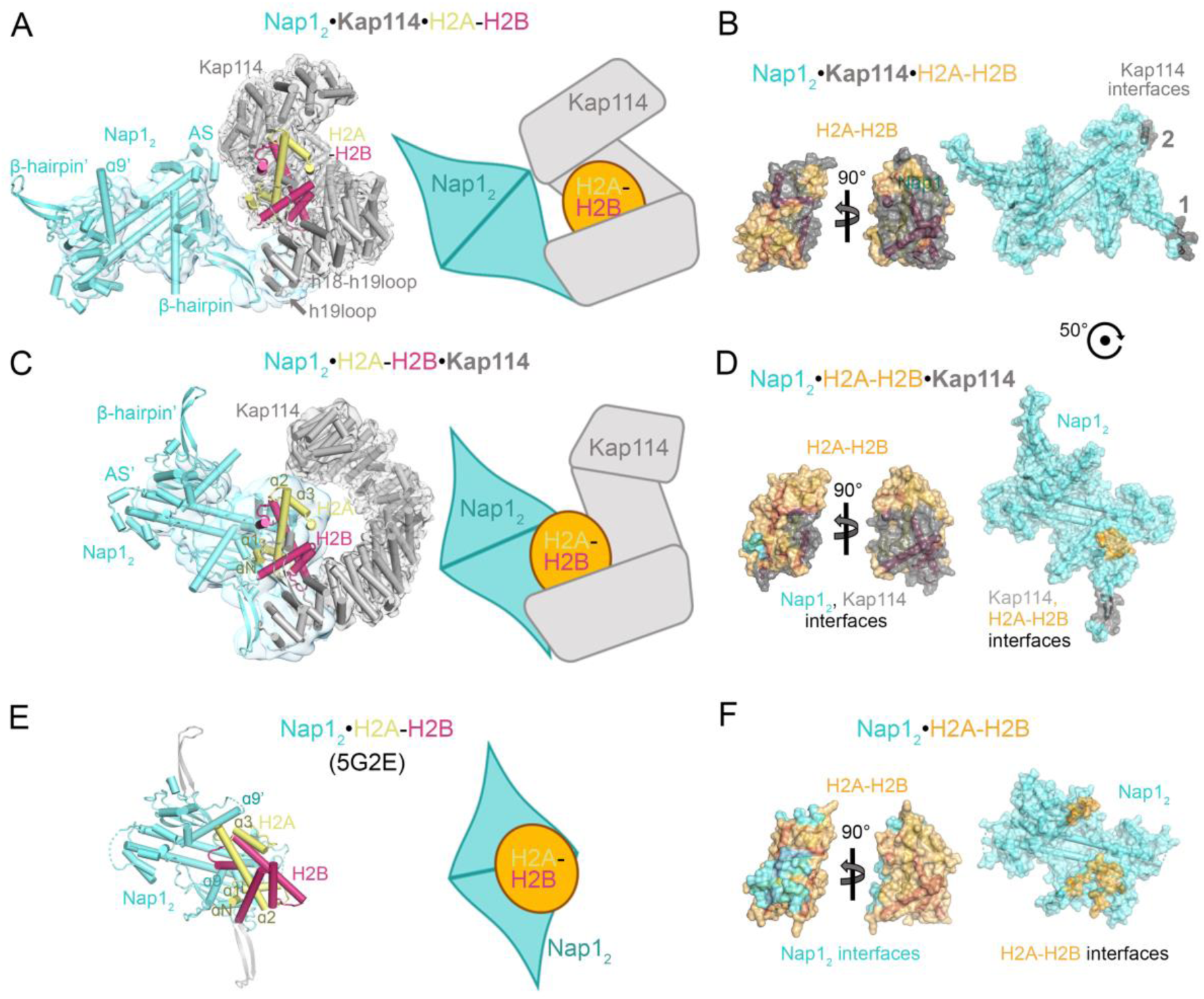
Cryo-EM structures reveal two distinct configurations of the Kap114/Nap1_2_/H2A-H2B complex. **(A)** Left, the Nap1_2_•**Kap114**•H2A-H2B (cyan•gray•yellow-pink) structure with overall/local refined maps (transparent gray/cyan surfaces). The β-hairpin and AS of the Nap1_2_ subunit contacting Kap114 are labeled while those of the other subunit are labeled β-hairpin’ and AS’. Right, a schematic of the structure. Note the similarity of Kap114 and H2A-H2B here with the binary Kap114•H2A-H2B complex in Figure 1B. **(B)** Nap1_2_•**Kap114**•H2A-H2B: H2A-H2B (orange; right view same as in **(A)**) and Nap1_2_ (cyan) surfaces, showing the Kap114 interfaces in gray. **(C)** Left, the Nap1_2_•H2A-H2B•**Kap114** structure with its consensus and local refined maps, aligned to H2A-H2B of **(A)**. Right, schematic. **(D)** Nap1_2_•H2A-H2B•**Kap114**: the histone (orange), Nap1_2_ (cyan) and Kap114 (gray) interfaces shown on Nap1_2_ and histone surfaces. The Nap1_2_ here is related to that in Nap1_2_•**Kap114**•H2A-H2B **(B)** by a 50°rotation. **(E)** The Nap1_2_•H2A-H2B (5G2E) crystal structure, with Nap1_2_ aligned to that in **(C)**. The Nap1_2_ β-hairpins here are disordered and here shown here in gray are hairpins from the structure in **(C)**; Nap1_2_ alignment in SI Appendix, Fig. S4E. Alignment of (**C)** and (**E**) in SI Appendix, Figs. S7 and 17. **(F)** Nap1_2_ and H2A-H2B interfaces of the 5G2E structure.

### Two ternary complex structures: the Nap1_2_•*Kap114*•H2A-H2B structure

One of the two ternary complex structures, Nap1_2_•**Kap114**•H2A-H2B, shows Kap114 wrapping around H2A-H2B while Nap1_2_ contacts only Kap114 (Fig. 2A and SI Appendix, Fig. S6A and B). The H2A-H2B-bound Kap114 here is very similar to the binary Kap114•H2A-H2B complex (Fig. 1B and SI Appendix, Fig. S7A); root mean square deviation (RMSD) of the two structures is ∼0.5 Å (41). The extensive Kap114-histone interactions in the two structures, that bury ∼2200 Å^2^ of H2A-H2B, are virtually the same (Figs. 1B, 2B and SI Appendix, Fig. S8).

Kap114 binds Nap1_2_ at two interfaces that involve the same Nap1_2_ subunit (Figs. 2B and 3A). Interface 1, formed by residues 931-942 in the acidic Kap114 h19loop binding to the basic β-hairpin of one Nap1_2_ subunit, covers 192 Å^2^ of Nap1_2_. Interface 2, formed by the basic Kap114 h6-h7loop being close to the negatively charged Nap1_2_ accessory subdomain (AS), covers 82 Å^2^ of Nap1_2_.

**Figure 3.**
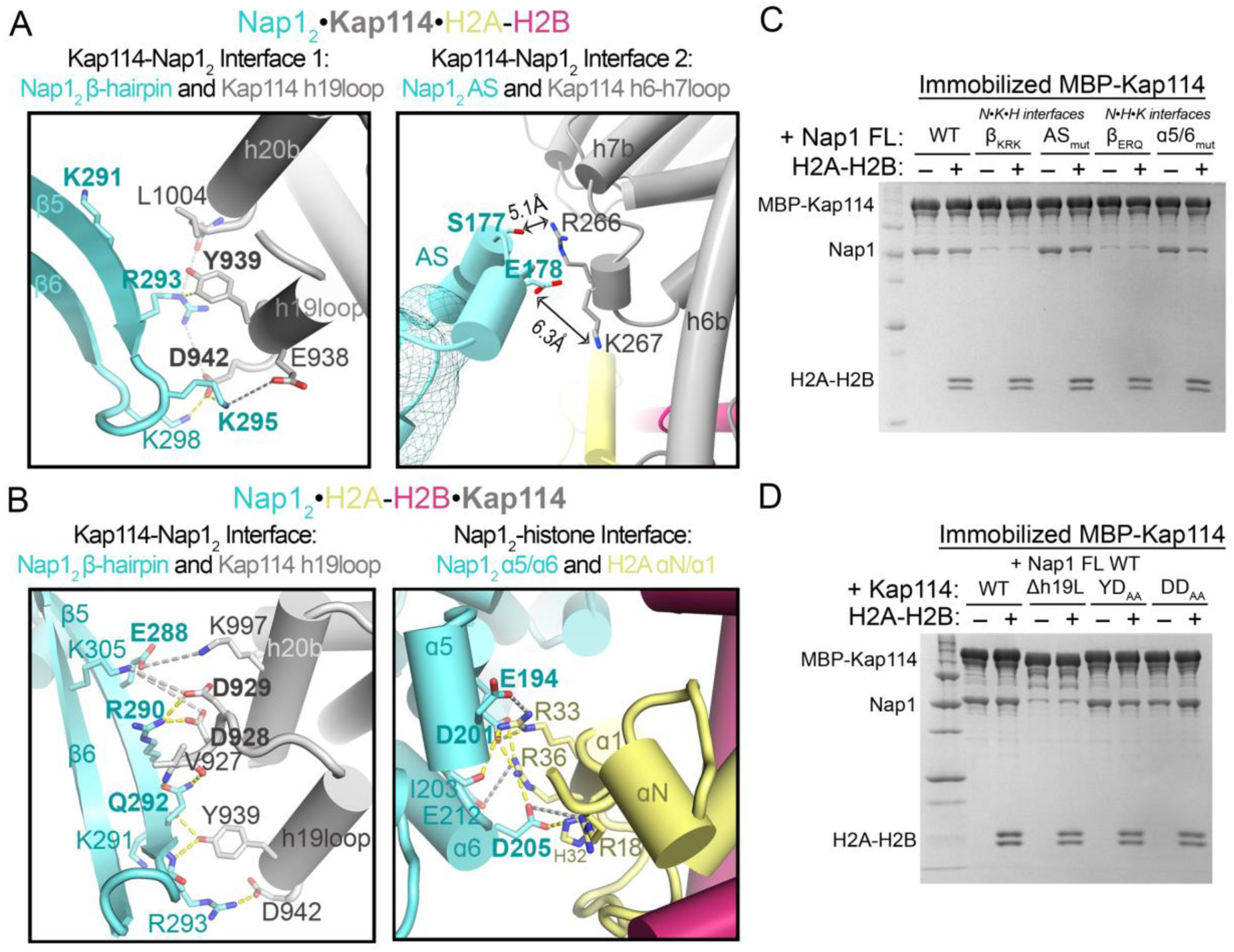
The two Kap114/Nap1_2_/H2A-H2B structures: interface details and mutagenesis. Dashed lines in **(A)** and **(B)** show intermolecular contacts <4.0 Å (yellow) and <8.0 Å (light grey). **(A)** Details of the Kap114 (grey)-Nap1_2_ (cyan) interfaces in the Nap1_2_•**Kap114**•H2A-H2B structure. Contacts are not drawn for Interface 2 because the Nap1_2_ AS is not observed in the final map. **(B)** Kap114-Nap1_2_ contacts (left) and H2A-H2B-Nap1_2_ contacts (right) in the Nap1_2_•H2A-H2B•**Kap114** structure. **(C)** Pull-down assay with 1 µM immobilized MBP-Kap114 and Nap1 FL WT or interface mutants β_KRK_ (K291A/R293A/K295A), AS_mut_ (S177A/E178A), β_ERQ_ (E288A/R290A/Q292A), α5/6_mut_ (E194A/D201A/D205A), ± equimolar H2A-H2B. Mutated residues are in bold in **(A)** and **(B)**. Both sets of Nap1_2_ β-hairpin mutations disrupted Kap114/Nap1_2_/H2A-H2B ternary complex formation. Neither Nap1_2_ AS nor histone-binding mutations had any effect. **(D)** Pull-down assay using 1 µM MBP-Kap114 WT or mutants Δh19L (h19loop deleted), YD_AA_ (Y939A/D942A) and DD_AA_ (D928A/D929A), with Nap1 FL ± H2A-H2B. Kap114 h19loop deletion disrupted ternary complex formation but double loop mutations did not. **(C)** and **(D)**, Coomassie-stained SDS-PAGE (controls in SI Appendix, Figs. S10).

### Two ternary complex structures: the Nap1_2_•H2A-H2B•*Kap114* structure

The Nap1_2_•H2A-H2B•**Kap114** structure is very different. Here, H2A-H2B is sandwiched between one subunit of the Nap1_2_ and the C-terminal HEAT repeats of Kap114 (Fig. 2C and SI Appendix, Fig. S6C and D). The 1555 Å^2^ interface of H2A-H2B with the C-terminal HEAT repeats of Kap114 is like the equivalent interfaces in Nap1_2_•**Kap114**•H2A-H2B and in other H2A-H2B-bound Kap114 structures (SI Appendix, Fig. S8). On the contrary, in the Nap1_2_•H2A-H2B•**Kap114** structure, the N-terminal half of Kap114 assumes a distinct, open conformation where repeats h1-h2 are ∼20 Å apart from the bound H2A-H2B, twice the distance observed in the structure of Ran^GTP^•Kap114•H2A-H2B (41) (SI Appendix, Fig. S7B).

Nap1_2_ in Nap1_2_•H2A-H2B•**Kap114** is tethered at two sites: one interface is with Kap114 and the other with H2A-H2B (Figs. 2D and 3B). The 271 Å^2^ Kap114-Nap1_2_ interface here involves the same Nap1_2_ β-hairpin and Kap114 h19loop that form Interface 1 of Nap1_2_•**Kap114**•H2A-H2B (Fig. 2B and D). However, Nap1_2_ in the two structures are oriented differently relative to the importins (Fig. 2A and C, and SI Appendix, Fig. S9). In Nap1_2_•**Kap114**•H2A-H2B, Nap1_2_ residues K291, R293 and K295, on one face of the β5 strand, contact the Kap114 h19loop (Fig. 3A). In Nap1_2_•H2A-H2B•**Kap114**, Nap1_2_ residues E288, R290 and Q292, on the opposite face of β5 strand, contact the Kap114 h19loop and the h20b helix (Fig. 3B).

The Nap1_2_-histone interface of Nap1_2_•H2A-H2B•**Kap114** involves acidic residues of the α4-6 helices of one Nap1_2_ subunit contacting basic residues of the H2A αN and α1 helix (Fig. 3B). This 207 Å^2^ interface is located at the histone interface in the binary Nap1_2_•H2A-H2B complex but is only ∼25% of the binary complex interface (Fig. 2D-F). However, the histone is far from exposed as more than 1500 Å^2^ of its surface is buried by Kap114 (Fig. 2E).

### Nap1_2_ β-hairpin is the hotspot for Kap114-binding

We mutated residues in both Kap114 and Nap1 that participate in Kap114-Nap1_2_ interactions to assess their importance in the two different Kap114/Nap1_2_/H2A-H2B structures. First, we mutated Nap1 residues in the two Kap114-Nap1_2_ interfaces of the Nap1_2_•**Kap114**•H2A-H2B structure (Fig. 3A). Alanine mutations of Nap1_2_ β-hairpin residues K291, R293 and K295 in Interface 1 (mutant β_KRK_) abolished pull-down by immobilized MBP-Kap114 in the absence and presence of H2A-H2B, while alanine mutations of the AS residues S177 and E178 in Interface 2 (mutant AS_mut_) did not (Fig. 3C). These results validated the Nap1_2_ β-hairpin as an energetically important Kap114-binding site and showed that the AS is less important for ternary complex formation.

Next, we mutated contact residues in the Kap114-Nap1_2_ and the Nap1_2_-H2A-H2B interfaces that are unique to the Nap1_2_•H2A-H2B•**Kap114** structure (Fig. 3B). Alanine mutations of the Nap1_2_ β-hairpin residues E288, R290 and Q292 (mutant β_ERQ_) that contact Kap114 abolished Nap1 pull-down by MBP-Kap114 in the absence and presence of H2A-H2B, while mutations of residues E194, D201 and D205 (mutant α5/6_mut_) that contact H2A-H2B did not (Fig. 3C). These results support the importance of the Nap1_2_ β-hairpin for Kap114-binding, in contrast to Nap1_2_-H2A-H2B interactions, which are less important for ternary complex formation.

We also mutated contact residues in Kap114. The Kap114 h19loop binds the Nap1_2_ β-hairpin in both ternary complex structures (Fig. 3A and B). It is therefore not surprising that truncating the Kap114 h19loop to remove all contact residues in both structures (mutant Kap114 Δh19L) abolished Kap114-Nap1 pull-down and formation of the ternary complex (Fig. 3D). However, mutating just Kap114 Y939 and D942 (mutant Kap114 YD_AA_) that contact Nap1_2_ solely in Nap1_2_•**Kap114**•H2A-H2B or mutating just D928 and D929 (mutant Kap114 DD_AA_) that contact Nap1_2_ solely in Nap1_2_•H2A-H2B•**Kap114**, did not affect Kap114-Nap1 pull-down (Fig. 3D). Many nearby acidic/electronegative Kap114 side chains may also interact with the Nap1_2_ β-hairpin (Fig. 3A and B).

In summary, mutagenesis findings indicate that although the interfaces between the Nap1_2_ β-hairpin and Kap114 h19loop are relatively small in both the Nap1_2_•Kap114•H2A-H2B and Nap1_2_•H2A-H2B•Kap114 structures, these interactions are essential for binding. Thus, these results validate the cryo-EM structures of both complexes. Our structural and biochemical results are also supported by previous studies that suggested the importance of the Nap1 b-hairpin for localization to the yeast nuclei (13, 36, 42).

### Nap1_2_•*Kap114*•H2A-H2B *versus* Nap1_2_•H2A-H2B•*Kap114*

The cryo-EM structures and mutagenic analysis above inform on how Kap114, Nap1_2_ and H2A-H2B may associate in the absence of Ran^GTP^, such as in the cytoplasm. Mutagenesis clearly shows that interactions between Nap1_2_ β-hairpin and Kap114 h19loop are critical for ternary complex formation even though cryo-EM analysis suggests the interactions are dynamic (Figs. 2 and 3). While challenging for structure determination, Nap1_2_ motions might influence interactions with its binding partners. Various Nap1_2_ structures suggest variations in protein dynamics. The cryo-EM map of unliganded Nap1_2_ has features for the entire homodimer (SI Appendix, Fig. S4A), contrasting with the crystal structure of the Nap1_2_•H2A-H2B complex where electron density for both sets of β-hairpins and AS regions was absent (Fig. 2E). In our ternary complex structures, the β-hairpins that bind Kap114 are resolved in the cryo-EM maps, whereas those of the other Nap1_2_ subunit are not (Fig. 2A and C). These structural observations may hint at dynamics-driven allostery and asymmetry within the conformationally symmetric Nap1_2_. Ligand binding, such as of histone and/or Kap114, does not alter Nap1_2_ conformation but may amplify its movements, resulting in an entropic penalty that prevents symmetric binding of the second ligand (43, 44). Increased Nap1_2_ motions upon Kap114 binding could contribute entropically to binding energy, stabilizing interactions, despite the small interfaces.

The interfaces between the Nap1_2_ β-hairpin and Kap114 differ in Nap1_2_•**Kap114**•H2A-H2B and Nap1_2_•H2A-H2B•**Kap114**. However, mutating either interface prevents the association of the three proteins. This implies that dynamic switching between the two configurations is essential for forming and maintaining the stability of the ternary complex. The Nap1_2_•**Kap114**•H2A-H2B and Nap1_2_•H2A-H2B•**Kap114** structures likely represent components within a dynamic ensemble of interconverting Kap114/Nap1_2_/H2A-H2B states in the cytoplasm. It is unclear which components of this ensemble will subsequently translocate through the NPC into the nucleus. The physical and chemical nature of the permeability barrier of the NPC and the mechanisms importin-cargo translocation across that barrier, including knowledge of importin-cargo conformations in the barrier, are far from understood at this time (45–48).

We need to consider another scenario. Our cryo-EM sample was prepared with Kap114, Nap1_2_ and excess H2A-H2B, which has been observed to destabilize the ternary Kap114/Nap1_2_/H2A-H2B complex (Fig. 1C-E). It is plausible that the structures of Nap1_2_•**Kap114**•H2A-H2B and/or Nap1_2_•H2A-H2B•**Kap114** structures could represent intermediates during disassembly of the ternary complex into binary Nap1_2_•H2A-H2B and Kap114•H2A-H2B complexes. Among these potential intermediates, the Nap1_2_•**Kap114**•H2A-H2B structure, characterized by a substantially smaller Nap1_2_ interface, might be a more favorable candidate for an intermediate state in the disassembly process.

Both Nap1_2_•**Kap114**•H2A-H2B and Nap1_2_•H2A-H2B•**Kap114** likely represent configurations of the Kap114/Nap1_2_/H2A-H2B ternary complex that could exist in the cytoplasm. However, their specific roles in the process of importing H2A-H2B into the nucleus are unclear and warrant deeper investigation. Understanding how Kap114/Nap1_2_/H2A-H2B interacts with Ran^GTP^, as is expected in the Ran^GTP^-rich nucleus, may provide insights into the distinct functions of the Nap1_2_•**Kap114**•H2A-H2B *versus* Nap1_2_•H2A-H2B•**Kap114** structures in the cytoplasmic millieu.

### Interactions between Kap114, Nap1 and H2A-H2B in the presence of Ran^GTP^

The Kap114/Nap1_2_/H2A-H2B ternary complex will encounter Ran^GTP^ upon entering the nucleus. All importins bind Ran^GTP^ tightly, almost always causing importin•cargo dissociation to release cargo in the nucleus (49). H2A-H2B import is an exception because Ran^GTP^ binds Kap114•H2A-H2B to form a stable ternary Ran^GTP^•Kap114•H2A-H2B complex (Fig. 4A) (41). Similarly, Kap114-Nap1 interaction persists in the presence of Ran^GTP^ as shown by pull-down assays and AUC analysis (Fig. 4B and SI Appendix, Fig. S12). SEC-MALS analysis showed that an equimolar mix of Kap114, Nap1_2_, H2A-H2B and Ran^GTP^ produced a major peak consistent with a complex that contains one molecule each of the four proteins (Fig. 4C).

**Figure 4.**
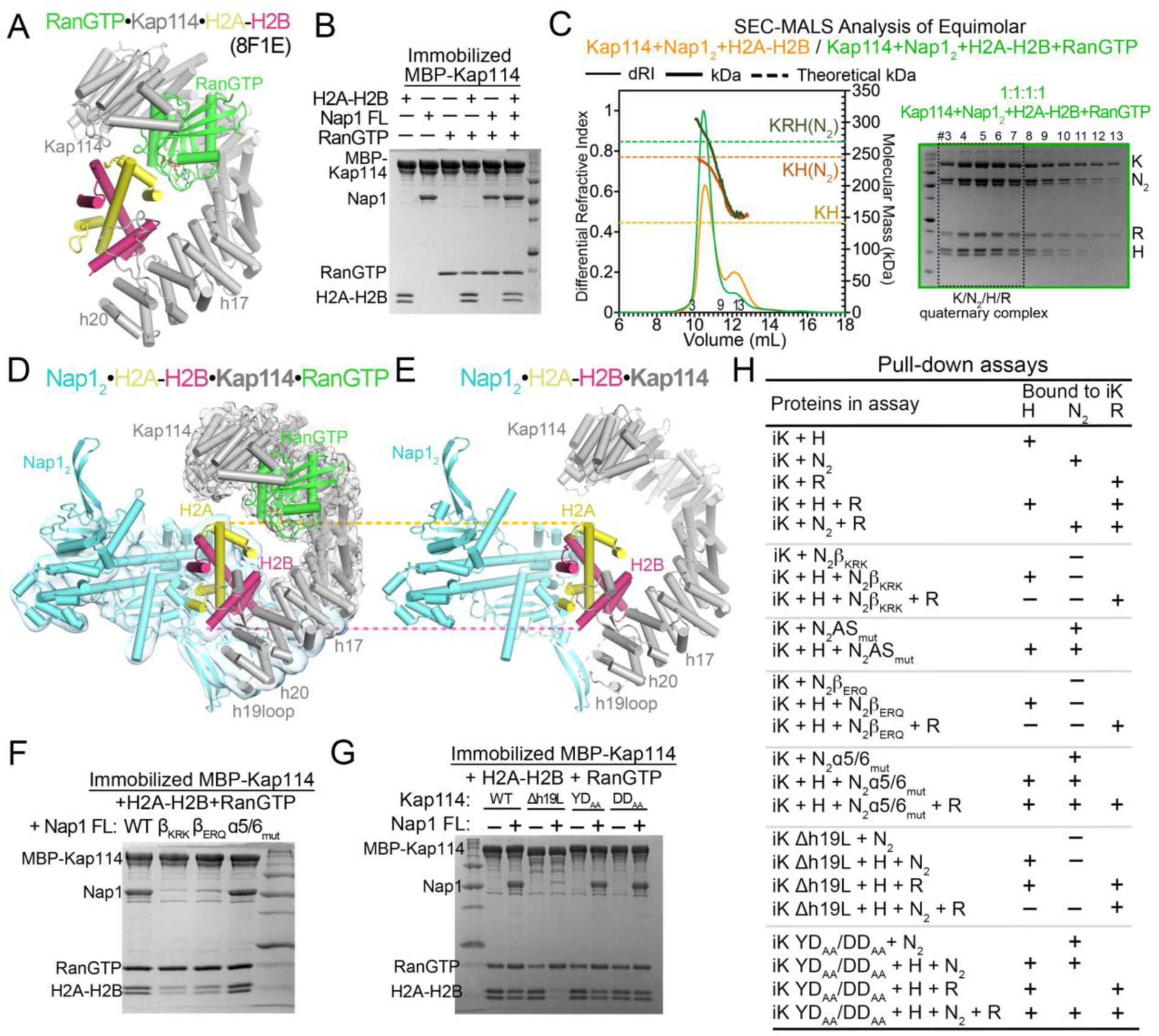
Structure of the Nap1_2_•H2A-H2B•Kap114•Ran^GTP^ complex. **(A)** The Ran^GTP^•Kap114•H2A-H2B structure (8F1E). **(B)** Pull-down assay with equimolar MBP-Kap114 (1 µM), Nap1_2_, H2A-H2B and Ran^GTP^; controls in SI Appendix, Fig. S12A-C. **(C)** SEC-MALS of Kap114, Nap1_2_, H2A-H2B and Ran^GTP^ (green curve), plotted like Fig. 1C, with gel of peak fractions. The major peak at 270-293 kDa has mass like the 1:1:1:1 Kap114/Nap1_2_/H2A-H2B/Ran^GTP^ complex (269 kDa). Kap114/Nap12/H2A-H2B SEC-MALS (orange, from Fig. 1D) is shown for comparison. Control in SI appendix, Fig. S12E. **(D)** The Nap1_2_•H2A-H2B•**Kap114**•Ran^GTP^ structure, with the consensus/local refined maps (gray/cyan) overlayed, shown next to the Nap1_2_•H2A-H2B•**Kap114** structure in **(E)**. **(A)** and **(D)** are aligned at Ran^GTP^, and **(D)** and **(E)** are aligned at H2A-H2B. Pull-down assays with **(F)** Nap1_2_ WT, β-hairpin mutants (β_KRK_ and β_ERQ_) and histone-binding site mutant α5/6_mut_, and **(G)** Kap114 h19loop mutants Δh19L, YD_AA_ and DD_AA_. Proteins in **(B)**, **(C)**, **(F)** and **(G)** visualized by Coomassie-stained SDS-PAGE. **(H)** Summary of all pull-down assays in Figs. 1, 3 and 4. iK refers to immobilized MBP-Kap114.

We solved the cryo-EM structure of this quaternary Kap114/Nap1_2_/H2A-H2B/Ran^GTP^ complex to 2.9 Å resolution (Fig. 4D and Si Appendix, Figs. S13 and S14 and Table S2). The final cryo-EM map is well-defined for Kap114 and Ran^GTP^ but the local resolution and map feature for Nap1_2_ and H2A-H2B are poor. Local refinement produced a 4.0 Å map that allowed docking of Nap1_2_ and H2A-H2B. We built the structure of the quaternary complex using Ran^GTP^ bound to the N-terminal HEAT repeats of Kap114 from the Ran^GTP^•Kap114•H2A-H2B structure (PDB: 8F1E) (41) and Nap1_2_, H2A-H2B and Kap114 repeats h17-h20 from the Nap1_2_•H2A-H2B•**Kap114** structure. We named this assembly the Nap1_2_•H2A-H2B•**Kap114**•Ran^GTP^ complex (Fig. 4D).

In the Nap1_2_•H2A-H2B•**Kap114**•Ran^GTP^ quaternary complex structure, Ran^GTP^ binds Kap114 the same as in the Ran^GTP^•Kap114•H2A-H2B structure (41) (Fig. 4A and D). Comparing Fig. 4D and E shows that Nap1_2_•H2A-H2B•**Kap114**•Ran^GTP^ is essentially Nap1_2_•H2A-H2B•**Kap114** with Ran^GTP^ bound at the N-terminal half of Kap114. Ran-binding in the quaternary complex stabilizes a Kap114 conformation where its N-terminus is closer to the bound H2A-H2B than in the ternary Nap1_2_•H2A-H2B•**Kap114** structure (Fig. 4D and E and SI Appendix, Fig. S7C). It is interesting that Nap1_2_•H2A-H2B•**Kap114**•Ran^GTP^ looks like Nap1_2_•H2A-H2B•**Kap114** rather than Nap1_2_•**Kap114**•H2A-H2B (compare Fig. 4D and E with Fig. 2A). This similarity may suggest that Nap1_2_•H2A-H2B•**Kap114** is the ternary configuration that enters the NPC for translocation into the nucleus.

Like the cytoplasmic Kap114/Nap1_2_/H2A-H2B ternary complex, mutation of the Nap1_2_ β-hairpin or truncation of the Kap114 h19loop abolished Kap114-Nap1_2_ pull-down in the presence of Ran^GTP^ and H2A-H2B (Fig. 4F-H), validating the Nap1_2_•H2A-H2B•**Kap114**•Ran^GTP^ cryo-EM structure and revealing that the Nap1_2_ β-hairpin remains the key Kap114-binding determinant in the presence of Ran^GTP^.

### Kap114 and Nap1_2_: co-chaperoning H2A-H2B in the absence of Ran^GTP^

Nap1 and Kap114/*Hs* IMP9 were previously identified as highly effective histone chaperones that prevent aggregation of H2A-H2B with DNA (13–16, 41, 50). Consistent with their histone chaperone function, the Nap1_2_ and Kap114 in both Nap1_2_•**Kap114**•H2A-H2B and Nap1_2_•H2A-H2B•**Kap114** shield substantial portions (∼1700-2200 Å^2^) of the nucleosomal H3-H4 and DNA binding regions of H2A-H2B (Fig. 2 and SI Appendix, Fig. S15).

We performed DNA competition assays to probe the chaperoning of H2A-H2B by Kap114 and Nap1_2_. In this assay, H2A-H2B chaperoning would result in disappearance of low-mobility DNA•H2A-H2B bands and appearance of high-mobility free DNA bands. We performed the experiments by either assembling DNA•H2A-H2B first and then titrating in Nap1_2_ and/or Kap114 (Fig. 5A) or assembling complexes of H2A-H2B with Kap114 and/or Nap1_2_ first before titrating them into DNA (SI Appendix, Fig. S16A). Regardless of the order of addition, Kap114 and Nap1_2_ together chaperone H2A-H2B from aggregation with DNA more effectively than either Kap114 or Nap1 alone.

**Figure 5.**
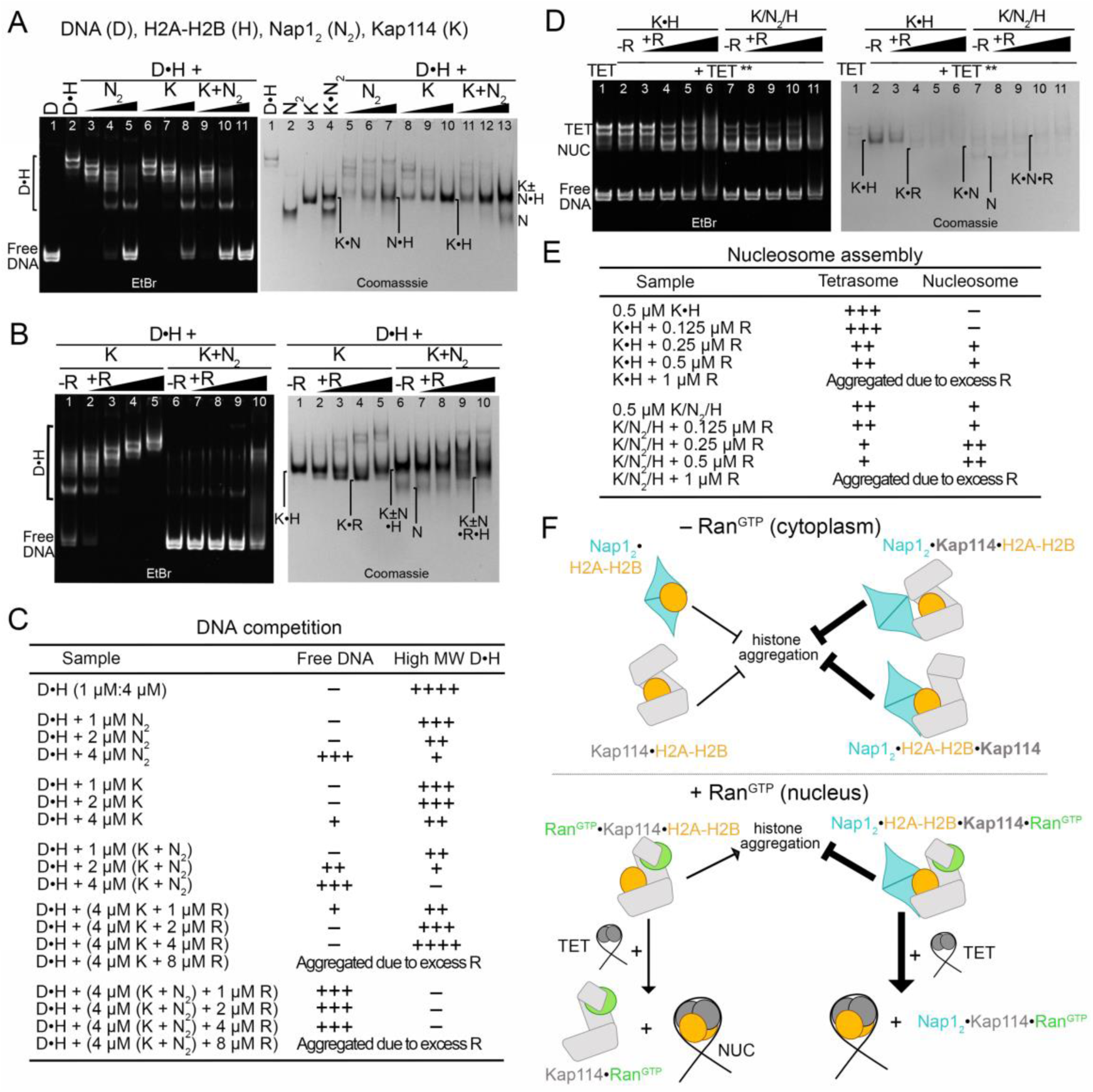
Kap114 and Nap1_2_ co-chaperone H2A-H2B in the presence of Ran^GTP^ for targeted release to tetrasomes. (A, B) DNA competition assays of DNA•H2A-H2B complex (D•H) added to 1-4 µM Nap1 (N_2_), Kap114 (K) or Kap114+Nap1_2_ mixture (K+N_2_), in the absence of Ran^GTP^ **(A)** or added to 4 µM K or K + N_2_ and 1-8 µM Ran^GTP^ (R) **(B)**. **(C)** Summary of DNA competition assay results. **(D)** Nucleosome assembly assay. Pre-assembled K•H or K/N_2_/H complexes without or with 0.125-1 µM R were added to tetrasomes (TET); **TET added last and nucleosome (NUC) formation tracked (details in methods). **(E)** Nucleosome assembly assay results. **(A, B** and **D)** visualized using ethidium bromide- and Coomassie-stained native PAGE gels. **(F)** Schematic of key findings: without Ran^GTP^ (e.g. cytoplasm), Kap114, Nap1_2_ and H2A-H2B assemble into a ternary complex that adopts at least two different configurations where Kap114 and Nap1_2_ co-chaperone H2A-H2B. In the presence of Ran^GTP^ (e.g. nucleus), Kap114 binds Ran^GTP^ to form the quaternary complex Nap1_2_•H2A-H2B•**Kap114**•Ran^GTP^ that inhibits histone aggregation and transfers H2A-H2B to assemble nucleosome (bottom right). Without Nap1_2_, H2A-H2B•Kap114•Ran^GTP^ does not inhibit histone aggregation (bottom left).

To summarize, when Ran^GTP^ is absent, as in the cytoplasm, Kap114 and Nap1_2_ within Kap114/Nap1_2_/H2A-H2B work together to enhance H2A-H2B sequestration compared to Kap114 alone. This effectively shields the histone from aberrant interactions.

### Kap114/Nap1 co-chaperoning and nucleosome assembly in the presence of Ran^GTP^

The quaternary Nap1_2_•H2A-H2B•**Kap114**•Ran^GTP^ structure buries 1847 Å^2^ of H2A-H2B. Here, Nap1_2_ and Kap114 together shield the DNA binding interfaces of H2A-H2B, reminiscent of their role in the cytosolic/ternary Nap1_2_•H2A-H2B•**Kap114** structure (Figs. 2D, 4D and SI Appendix, Fig. S15). The similarity in their protein arrangements suggests that, even in the presence of Ran^GTP^, Kap114 and Nap1_2_ are likely to prevent H2A-H2B from aggregating with DNA. We tested this hypothesis with DNA competition assays using Kap114, H2A-H2B, with/without Nap1_2_ and titrating in Ran^GTP^ (Fig. 5B and C, SI Appendix, Fig. S16B). When Ran^GTP^ is absent, Kap114 alone effectively chaperones H2A-H2B; however, its efficacy as a histone chaperone decreases substantially in the presence of the GTPase. Conversely, when both Nap1_2_ and Kap114 are present, they efficiently prevent non-specifically aggregation of H2A-H2B with DNA irrespective of whether Ran^GTP^ is present or absent. Therefore, within the quaternary/nuclear Nap1_2_•H2A-H2B•**Kap114**•Ran^GTP^ complex, Kap114 and Nap1_2_ cooperate to successfully chaperone H2A-H2B. SI Appendix Fig. S15 compares H2A-H2B surfaces in various complexes.

We performed nucleosome assembly assays to assess the effects of Kap114, Nap1_2_ and Ran^GTP^ on depositing H2A-H2B to form nucleosomes (Fig. 5D, E and SI Appendix, Fig. S16). As shown previously, Kap114 alone does not deposit H2A-H2B onto tetrasomes (41), but Ran^GTP^ facilitates efficient histone transfer from Kap114 to form nucleosomes. Nap1_2_ facilitates efficient histone transfer from Kap114 to form nucleosomes when Ran^GTP^ absent. However, nucleosome assembly is most efficient when Kap114, Nap1_2_ and Ran^GTP^ are all present, as evidenced by the almost complete disappearance of tetrasomes in lanes 9 – 11 of Fig. 5D.

In summary, Kap114 and Nap1_2_ in Nap1_2_•H2A-H2B•**Kap114**•Ran^GTP^ cooperate to chaperone H2A-H2B (Fig. 5F). Without Nap1_2_, Ran^GTP^ would bind to the Kap114•H2A-H2B complex to reorient the bound histone and increase its exposure to DNA. However, Nap1_2_ binding H2A-H2B alongside Kap114 in the nuclear/quaternary Nap1_2_•H2A-H2B•**Kap114**•Ran^GTP^ complex shields the histone from DNA and positions it for effective transfer into nucleosomes (Fig. 5F).

### H2A-H2B transfer from Nap1_2_•H2A-H2B•Kap114•Ran^GTP^ to nucleosome

We used a Kap114 mutant that cannot bind Nap1_2_ to understand how Kap114, Nap1_2_ and Ran^GTP^ might work together to transfer H2A-H2B onto the assembling nucleosome. Pull-down binding assays were conducted with equimolar Kap114 Δh19L, Nap1_2_, H2A-H2B and Ran^GTP^ (Figs. 3D, 4G and H). The Kap114 Δh19L mutant, unable to bind Nap1_2_, still managed to capture H2A-H2B in the presence of Nap1_2_, forming the Kap114 Δh19L•H2A-H2B complex (Figs. 3D and 4H), consistent with H2A-H2B’s high affinity for the importin (41). Kap114 Δh19L also pulled down H2A-H2B in the presence of Ran^GTP^ to form Ran^GTP^•Kap114 Δh19L•H2A-H2B (Fig. 4G and H). However, Kap114 Δh19L failed to capture H2A-H2B in the presence of both Ran^GTP^ and Nap1_2_, suggesting that H2A-H2B may be binding to unbound Nap1 instead (Fig. 4G and H). When Ran^GTP^ was present, free Nap1, unable to bind Kap114 Δh19L, effectively displaced histones from the importin. These results imply that Ran^GTP^ in the Nap1_2_•H2A-H2B•**Kap114**•Ran^GTP^ complex likely promotes the transfer of H2A-H2B from Kap114 to Nap1_2_, facilitating optimal deposition of H2A-H2B into nucleosomes. The quaternary assembly may provide a path to specifically move H2A-H2B from Kap114 to Nap1_2_ to the tetrasome without losing the histone to non-specific/aberrant interactions in the nucleus.

## Conclusion

In an environment devoid of Ran^GTP^, such as in the cytoplasm, equimolar Kap114, Nap1_2_ and H2A-H2B assemble into a 1:1:1 Kap114/Nap1_2_/H2A-H2B ternary complex. Two cryo-EM structures, Nap1_2_•**Kap114**•H2A-H2B and Nap1_2_•H2A-H2B•**Kap114**, revealed two distinct configurations of the three proteins, potentially depicting various states within a dynamic equilibrium of Kap114/Nap1_2_/H2A-H2B complexes in the cytoplasm. The DNA-binding surfaces of H2A-H2B in both structures are shielded and DNA competition assays showed that Kap114 and Nap1_2_ together co-chaperone the histone from aberrant interactions more effectively than either protein alone. The binding of Ran^GTP^, simulating condition within the nucleus, results in the Nap1_2_•H2A-H2B•**Kap114**•Ran^GTP^ quaternary complex, which assumes the protein arrangement observed in the Nap1_2_•H2A-H2B•**Kap114** ternary structure, with Ran^GTP^ binding to the N-terminal HEAT repeats of Kap114. The Nap1_2_•H2A-H2B•**Kap114**•Ran^GTP^ complex relies on Nap1_2_’s critical role for effective H2A-H2B chaperoning in the presence of Ran^GTP^. This complex efficiently deposits H2A-H2B to form nucleosomes. Through delineating the biochemical mechanism of Nap1_2_ as a co-factor in H2A-H2B nuclear import, together with Kap114, this study explains how histones H2A-H2B are co-chaperoned and seamlessly transferred into assembling nucleosomes.

## Materials and Methods

### Pull-down binding Assays

Details for protein constructs, expression, and purification are in SI methods. All biochemical and biophysical studies were conducted at an ionic strength close to physiological salt levels of 150 mM NaCl. *In vitro* pull-down binding assays were performed in triplicate by incubating proteins at their indicated concentrations with 20 µL amylose resin (bead bed volume; New England Biolabs) in a 200 µL reaction in Assay Buffer (20 mM HEPES pH 7.4, 150 mM NaCl, 2 mM MgCl2, 10% (v/v) glycerol and 2 mM DTT) for 30 min at 4°C, followed by washing with 500 µL buffer twice. Bound proteins were separated and visualized by SDS-PAGE and Coomassie Blue staining. Nap1 band intensities were quantified using Bio Rad Image Lab software, adjusted for the intensity of the MBP-Kap114 band and normalized to Nap1 WT control.

### Analytical Ultracentrifugation

Individual proteins were dialyzed overnight into AUC buffer containing 20 mM Tris-HCl pH 7.5, 150 mM NaCl, 2 mM MgCl_2_ and 2 mM TCEP, at 4°C and assembled in the AUC sample chamber at the indicated concentrations in 400 µL. Sedimentation coefficients were measured by monitoring absorbance at 280 nm in a Beckman-Coulter Optima XL-1 Analytical Ultracentrifuge. Time stamps were corrected using REDATE (51). SEDNTERP was used to calculate the buffer density, buffer viscosity, and protein partial-specific volumes (52). SEDFIT was used to calculate sedimentation coefficient distributions c(*s*) where the regularization calculated a confidence level of 0.68 was used, time-independent noise elements were accounted for, and at a resolution of 100 (53). SEDFIT was also used to estimate molecular weight by obtaining the sedimentation coefficient through integration of c(*s*) and the frictional ratios by refining the fit of the model. The c(*s*) distribution and isotherm integration was done with GUSSI (54). Further isotherm fitting was performed using SEDPHAT (55).

### Size Exclusion Chromatography Multi-Angle Light Scattering

SEC-MALS experiments were performed in Assay buffer without glycerol with proteins at the indicated molar ratios to 80 µM Kap114 following established protocol, including sample dialysis and buffer filtration (56). Concentration of proteins are high in the small injection of 100 µL to ensure complete complex formation when it is diluted in the SEC column. The data was collected and processed using ASTRA software using default settings with no de-spiking (Wyatt Technology). The run for 80 µM Kap114 + 80 µM H2A-H2B + 80 µM Nap1_2_ was interrupted at the void volume due to fractionation issues causing a total pause of 1.895 minutes. A correction of 1.38933 mL was applied to match other runs.

### Cryo-EM Sample preparation and data collection and analysis

To assemble the Kap114/H2A-H2B/Nap1 complex, *Sc* H2A-H2B, Nap1 core, and Kap114 were dialyzed overnight into cryo-EM buffer containing 20 mM Tris pH 7.5, 150 mM NaCl, and 2 mM DTT. A 1:2 molar ratio mixture of Nap1 core dimer:H2A-H2B was incubated at room temperature for 10 minutes followed by addition of 1 molar ratio of Kap114, followed by rapid addition of glutaraldehyde to a final concentration of 0.05%. Crosslinking proceeded for one minute before quenching and removal of glutaraldehyde by SEC using a Superdex200 10/300 Increase column (Cytiva) that was equilibrated with cryo-EM buffer containing TCEP (instead of DTT). 0.5 mL fractions were collected for mass photometry (MP) analysis (details in SI Methods). To assemble the Ran^GTP^/Kap114/Nap1/H2A-H2B complex, a 10 mg/mL mix of a stoichiometric molar ratio of 1 Kap114 to 1 H2A-H2B to 2 Nap1_2_ FL (His tag intact) to 1 Ran^GTP^ was dialyzed overnight before crosslinking with 0.05% glutaraldehyde for 1 minute and separation by SEC in Assay buffer without glycerol. Fractions of 0.25 mL were collected for MP analysis. Fractions with the least aggregated species and most enriched with the relevant complexes (described in SI Appendix, Figs. S3A and S13A) were selected for grid preparation.

For Kap114/H2A-H2B/Nap1, the sample was diluted to ∼1.2 mg/mL and supplemented with 0.003125% (w/v) Triton X-100. For Ran^GTP^/Kap114/Nap1/H2A-H2B, sample was diluted to a ∼1.5 mg/mL with 0.003125% (w/v) Tyloxapol. Quantifoil grids were glow-discharged, blotted, and plunge-frozen. Grids were screened and data was collected and analyzed as described in SI Methods.

### DNA competition assays

601 Widom DNA was purified from pUC19-30X601 grown in C2925 dam^-^dcm^-^ cells (gift from Dr. Mike Rosen) using Qiagen Giga prep kit following manufacturer’s protocol. The DNA was resuspended in 10 mM Tris pH 7.4, 1 mM EDTA in the last step of purification. EcoRV (New England Biolabs) cleavage was performed in CutSmart buffer overnight, and the DNA extracted using PCIAA and precipitated in sodium acetate pH 5.2 and ethanol. The DNA was then rinsed with ethanol, dried, and resolubilized in 10 mM Tris pH7.4, 10 mM MgCl_2_. The cut insert was separated from the vector using PEG8000 precipitation and subjected to a final ion exchange purification on a DEAE-FF column before it was concentrated and frozen.

DNA was mixed with H2A-H2B, Kap114 and/or Nap1_2_ FL with and without Ran^GTP^ at the indicated concentrations in 15 µL reactions in Assay Buffer. DNA was either pre-mixed with H2A-H2B in a 3X concentration master-mix, or alternatively, added at the end. Reactions were incubated at room temperature for 30 min and then 5 µL of sample buffer containing 20% (v/v) glycerol and bromophenol blue was added. 8 µL each of this gel sample was ran on two separate Native 5% PAGE gels with 0.5X TBE at 150V for 40 min. One gel was stained with EtBr in water and the other with Coomassie Blue. Assays were performed in duplicates.

### Nucleosome assembly assays

Tetrasomes were made by mixing *Xl* H3-H4 tetramers and DNA at 1:1 ratio (DNA conc = 0.7 mg/mL) in 10 mM HEPES pH7.4, 1 mM EDTA, 2 M NaCl, 1 mM DTT and then dialyzed step-wise from 2 M NaCl (0.5 hr) to 1M (1 hr), 0.85 M (1.5 hr), 0.65 M (1.5 hr), 0.2 M (1.5 hr) and 0.15 NaCl M (overnight) at 4°C. The volume was measured at the end and reaction was diluted to 2 µM with Assay buffer to 2X working concentration immediately before use, assuming 100% formation of tetrasomes. 12 µL reactions with *Sc* H2A-H2B, Kap114, Nap1_2_ FL and tetrasomes with or without Ran^GTP^ were assembled at indicated concentrations in figure legends of Figure 5 and incubated together at room temperature for 1 hr before addition of 4 µL of sample buffer. 7 µL of the resulting samples were visualized the same way as DNA competition assays. Assays were performed in duplicates.

The input sample in lane 1 of Fig. 5D shows that tetrasome formation is not 100% complete and some free DNA was present. Free DNA was present in all reactions and not incorporated into nucleosomes, suggesting that any free H3-H4 not incorporated into tetrasomes was inactive/degraded. The control experiment without Ran^GTP^ (SI Appendix, Fig. S16C) shows that no tetrasomes remain after addition of 1 µM H2A-H2B, suggesting that the tetrasome concentration was likely < 0.5 µM. This is consistent with the results of lane 10 Fig. 5D where nucleosome formation is not 100% complete with the addition of just 0.5 µM H2A-H2B. The concentration of 0.5 µM H2A-H2B used in the experiment shown in Fig 5D was selected to capture differences in the effectiveness of the Kap114/Nap1_2_/H2A-H2B ternary complex and the binary Kap114•H2A-H2B complex to deposit H2A-H2B onto tetrasomes, in the presence and absence of Ran^GTP^.

## Acknowledgments

We thank all members of the Structural Biology Laboratory (SBL) and the CEMF in UTSW for their expert assistance with cryo-EM data collection. We greatly appreciate Macromolecular Biophysics Resource (MBR) and Chad Brautigam for discussion and use of the AUC and Mass Photometer. We also thank the Erzberger lab for using their plate reader for FP assays. We thank Rosen Lab for help with the Widom DNA purification. We also thank Casey Wing, Mike Rosen and Mike Rout for critical reading of the manuscript. SBL and CEMF at UTSW are partially supported by grant RP220582 from the Cancer Prevention & Research Institute of Texas (CPRIT). The Refeyn Mass Photometer in MBR was supported by grant S10-OD030312 from the National Institute of Health. We thank the D’Arcy lab for the use of their SEC-MALS equipment and Joy Shaffer for training. Molecular graphics and analyses performed with UCSF ChimeraX, developed by the Resource for Biocomputing, Visualization, and Informatics at the University of California, San Francisco, with support from National Institutes of Health R01-GM129325 and the Office of Cyber Infrastructure and Computational Biology, National Institute of Allergy and Infectious Diseases. This research used resources of the Advanced Photon Source, a U.S. Department of Energy (DOE) Office of Science user facility operated for the DOE Office of Science by Argonne National Laboratory under Contract No. DE-AC02-06CH11357. This work was funded by NIGMS of NIH under Awards R35GM141461 (Y.M.C.), R01GM069909 (Y.M.C.), T32GM008203 (J.J), the Welch Foundation Grants I-1532 (Y.M.C.), support from the Alfred and Mabel Gilman Chair in Molecular Pharmacology, Eugene McDermott Scholar in Biomedical Research (Y.M.C.) and the Gilman Special Opportunities Award (H.Y.J.F.).

