## Supplementary Information for "Nap1 and Kap114 co-chaperone H2A-H2B and facilitate targeted histone release in the nucleus"

**Table S1. Cryo-EM data collection, refinement, and validation statistics.**

| | **Nap1$_2$** | **Nap1$_2$•*Kap114*•H2A-H2B** | | | **Nap1$_2$•H2A-H2B•*Kap114*** | | |
|---|---|---|---|---|---|---|---|
| | PDB: 9B23 EMD-44095 | Consen-sus Map EMD-44122 | Locally refined map for Nap1$_2$ EMD-44121 | Composite Map PDB: 9B31 EMD-44120 | Consensus Map EMD-44140 | Locally refined map for Nap1$_2$ •H2A-H2B EMD-44137 | Composite Map PDB: 9B3F EMD-44136 |
| **Data collection and processing** | | | | | | | |
| Magnification (X) | 105,000 | | | | | | |
| Voltage (kV) | 300 | | | | | | |
| Electron exposure (e$^-$/Å$^2$) | 52 | | | | | | |
| Defocus range (μm) | 1.5-2.5 | | | | | | |
| Pixel size | 0.83 | | | | | | |
| Symmetry (Å) | C1 | | | | | | |
| Initial particle images | 4,314,112 | | | | | | |
| Final particle no. | 230,210 | 148,410 | | | 113,011 | | |
| Map resolution (Å) | 3.21 | 3.20 | 4.84 | | 3.54 | 5.62 | |
| FSC threshold | 0.143 | | | | | | |
| **Refinement** | | | | | | | |
| Initial model used (PDB code) | AlphaFold-multimer Nap1 FL model | | | AF-P53067-F1, 8F0X, 9B23 | | | AF-P53067-F1, 8F1E, 9B23 |
| Model composition | | | | | | | |
| Non-H Atoms | 4,670 | | | 13,559 | | | 13,451 |
| Protein residues | 568 | | | 1,682 | | | 1,668 |
| Mean *B* factors (Å$^2$) | 196.99 | | | 129.6 | | | 317.38 |
| R.m.s. deviations | | | | | | | |
| Bond lengths (Å) | 0.003 | | | 0.004 | | | 0.002 |
| Bond angles (°) | 0.592 | | | 0.543 | | | 0.505 |
| CCvolume/mask | 0.68/0.69 | | | 0.76/0.75 | | | 0.70/0.70 |
| **Validation** | | | | | | | |
| MolProbity score | 1.69 | | | 1.30 | | | 1.46 |
| Clashscore | 10.54 | | | 5.54 | | | 7.64 |
| Poor rotamers (%) | 0 | | | 0 | | | 0 |
| Ramachandran plot | | | | | | | |
| Favored (%) | 97.16 | | | 98.26 | | | 97.82 |
| Allowed (%) | 2.84 | | | 1.74 | | | 2.18 |
| Disallowed (%) | 0 | | | 0 | | | 0 |
| CaBLAM outliers (%) | 1.25 | | | 1.15 | | | 1.04 |
| EMRinger score | 0.94 | | | 1.87 | | | 0.94 |

**Table S2. Cryo-EM data collection, refinement, and validation statistics continued.**

| | $Nap1_2$•H2A-H2B•*Kap114*•$Ran^{GTP}$ | | |
|---|---|---|---|
| | Consensus Map<br>EMD-44151 | Locally refined map for $Nap1_2$•H2A-H2B<br>EMD-44150 | Composite Map<br>PDB: 9B3I<br>EMD-44141 |
| **Data collection and processing** | | | |
| Magnification | 165,000X | | |
| Voltage (kV) | 300 | | |
| Electron exposure ($e^-/Å^2$) | 50 | | |
| Defocus range (μm) | 0.9-2.2 | | |
| Pixel size (Å) | 0.738 | | |
| Symmetry imposed | C1 | | |
| Initial particle images (no.) | 1,381,753 | | |
| Final particle images (no.) | 133,516 | | |
| Map resolution (Å) | 2.88 | 3.97 | |
| FSC threshold | 0.143 | | |
| **Refinement** | | | |
| Initial model used (PDB code) | | | 8F1E, 9B3F |
| Model composition | | | |
| Non-hydrogen Atoms | | | 14,951 |
| Protein residues | | | 1,854 |
| Mean *B* factors ($Å^2$)<br>Protein/Ligand | | | 112.36/40.96 |
| R.m.s. deviations | | | |
| Bond lengths (Å) | | | 0.006 |
| Bond angles (°) | | | 0.781 |
| CCvolume/mask | | | 0.74/0.73 |
| **Validation** | | | |
| MolProbity score | | | 1.55 |
| Clashscore | | | 10.80 |
| Poor rotamers (%) | | | 0.24 |
| Ramachandran plot | | | |
| Favored (%) | | | 98.20 |
| Allowed (%) | | | 1.80 |
| Disallowed (%) | | | 0 |
| CaBLAM outliers (%) | | | 0.82 |
| EMRinger score | | | 2.45 |

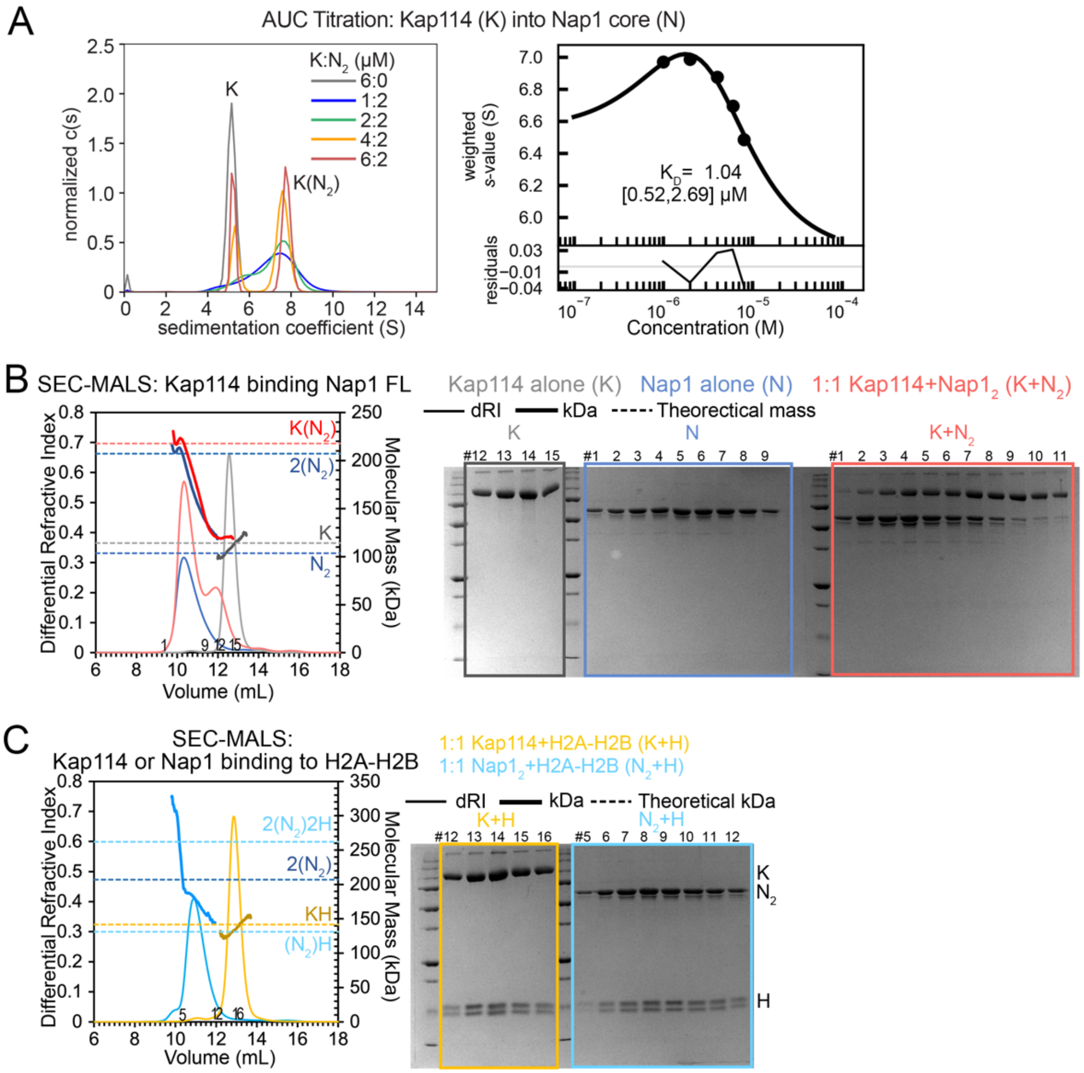


**Figure S1. Binary interactions between Kap114, Nap1 and H2A-H2B. (A)** AUC (left) and binding isotherm (right) of Kap114 (K; 5.2 S) titration into the Nap1 core dimer ($N_2$) at the concentrations indicated. Molecular weight estimate, using the c(s) distribution of the most saturated 6:2 sample, of the 7.9 S complex was 178 kDa, consistent with a K•$N_2$ complex (theoretical molecular weight, 186 kDa). The isotherm was generated using a one-site binding model and fitting residuals are plotted below. The dissociation constant or $K_D$ is shown with the values in brackets representing 95% confidence interval. **(B)** SEC-MALS analysis of Nap1 FL (N; blue), Kap114 (K; gray) and 1:1 mixture of both (red; 80 μM in 100 μL injection sample). The differential refractive index (dRI) traces are plotted as thin lines (left y-axis) and the molecular mass (kDa) traces as thick lines (right y-axis). Theoretical masses of the indicated proteins are marked with dashed lines. Peak fractions were visualized by Coomassie-stained SDS-PAGE (right). N formed tetramers with apparent molecular mass of ~200 kDa (elution volume ~10.3 mL), consistent with previous reports (1, 2). K eluted at ~12.5 mL with expected apparent molecular mass ~110 kDa. A 1:1 molar mixture of K and $N_2$ formed a peak of ~220 kDa that matches a K•$N_2$ complex. The increase in DRI signal of the K+$N_2$ compared to the N traces is consistent with incorporation of one K molecule. **(C)** SEC-MALS experiment for Kap114 (K; yellow) or Nap1 FL (N; cyan) binding to H2A-H2B (H) at the indicated molar ratios, plotted as in **(B)**. K+H eluted at the expected ~13 mL with the expected molecular mass whereas the $N_2$+H mixture eluted at volumes that span molecular masses between those that match the 2$N_2$•2H complex and the $N_2$•H complex, indicating that one or two copies of H was incorporated into mixture of $N_2$ and 2$N_2$, respectively.

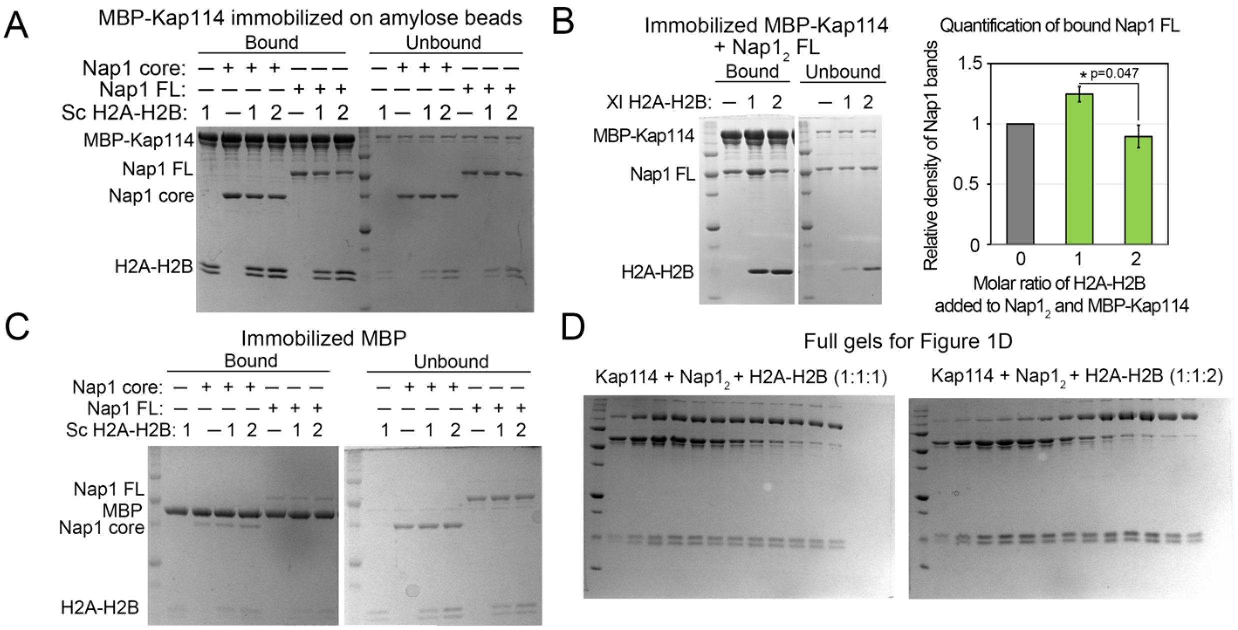


**Figure S2. Interactions between Kap114, Nap1 and H2A-H2B. (A)** The full gel of the binding assay shown in **Figure 1C**: 1 μM immobilized MBP-Kap114 and $Nap1_2$ core of FL with or without 1 or 2 μM *Sc* H2A-H2B. Bound and unbound proteins after extensive washing were visualized by Coomassie-stained SDS-PAGE. **(B)** Pull-down binding assay as in **A**, but with Nap1 FL and *Xl* H2A-H2B. Quantification of the average relative intensities of triplicate experiments of the bound FL Nap1, when normalized to the sample without H2A-H2B, is plotted with error bars that indicate standard deviation (s.d.). Like chicken H2A-H2B (3), but unlike *Sc* H2A-H2B, *X. laevis* (*Xl*) H2A-H2B increased *Sc* Nap1 association with Kap114, suggesting that different H2A-H2B homologs bind *Sc* Nap1 and Kap114 differently. Student t-test shows significant difference between 1 and 2 μM H2A-H2B samples, where less Nap1 was pulled down in the presence of excess H2A-H2B, suggesting destabilization of the ternary Kap114/$Nap1_2$/H2A-H2B complex. **(C)** Control pull-down experiment of 1 μM MBP (immobilized) and equimolar $Nap1_2$ without or with H2A-H2B (1 or 2 molar ratio). Background binding of Nap1 to the immobilized MBP was minimal. **(C)** Full gels for the gel images in **Figure 1D**.

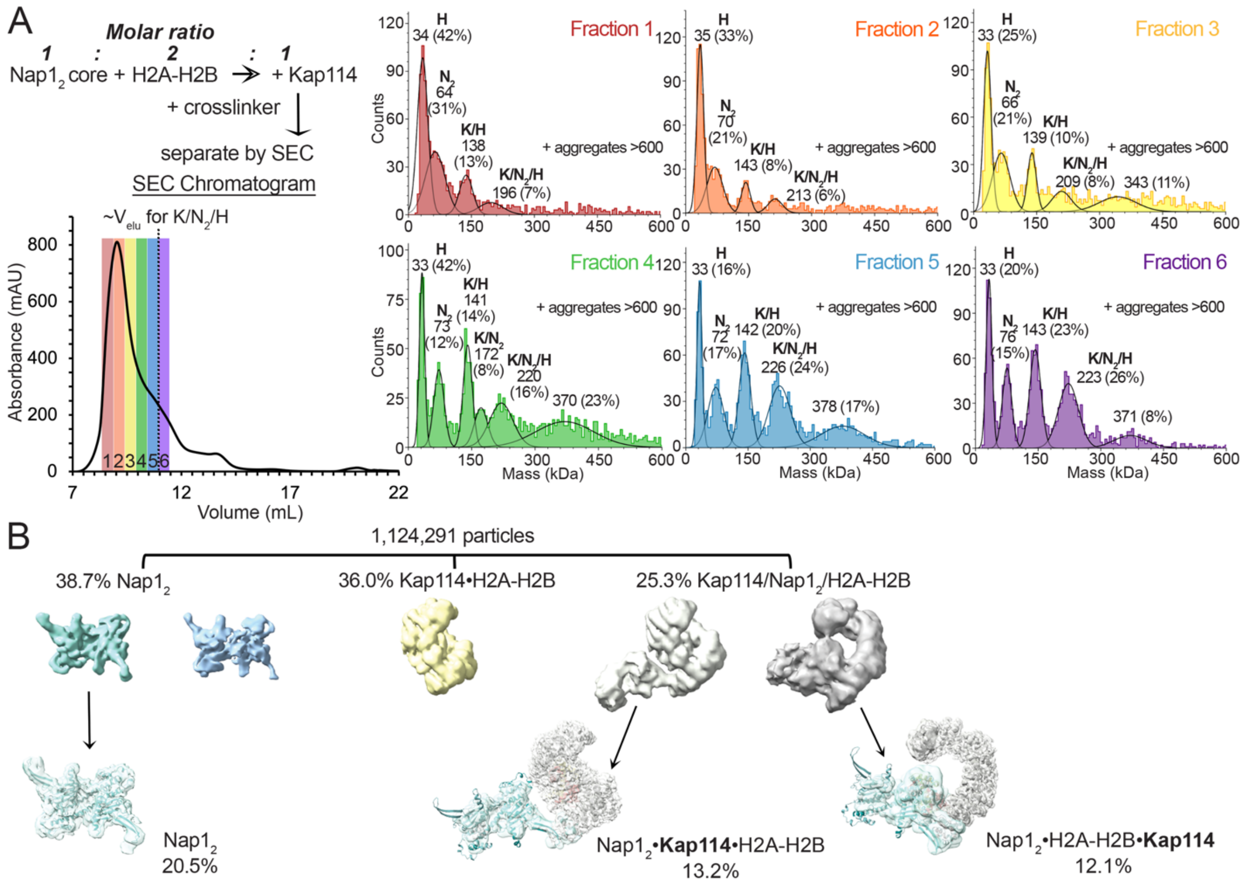


**Figure S3. Cryo-EM sample heterogeneity of crosslinked complex of Kap114 with Nap1 core and H2A-H2B. (A)** Top left, the scheme for assembly and crosslinking of mixture of Kap114 (K), Nap1 (core) dimer ($N_2$) and H2A-H2B (H), followed by size-exclusion chromatography (SEC). Bottom left, the SEC chromatogram with fractions 1-6 (red to purple) that were then subjected to mass photometry. The typical elution volume of an un-crosslinked complex of all three proteins is ~11 mL (dotted line). Mass photometry data of the six SEC fractions are plotted on the right. Aggregates beyond 600 kDa were not displayed. The mean masses (kDa) and relative populations (%) are indicated above the fitted gaussian peaks, along with the likely protein/complex that corresponds to the approximate mass. The species of ~340-380 kDa may be crosslinked complexes of K/$N_2$/H with another copy of K/H; such large complex was not observed in AUC or SEC-MALS studies. Fraction 6, most enriched with the complex containing all three proteins in a 1:1:1 ratio and has the least large aggregates, was used for cryo-EM grid preparation. **(B)** Particle distribution of Cryo-EM data obtained.

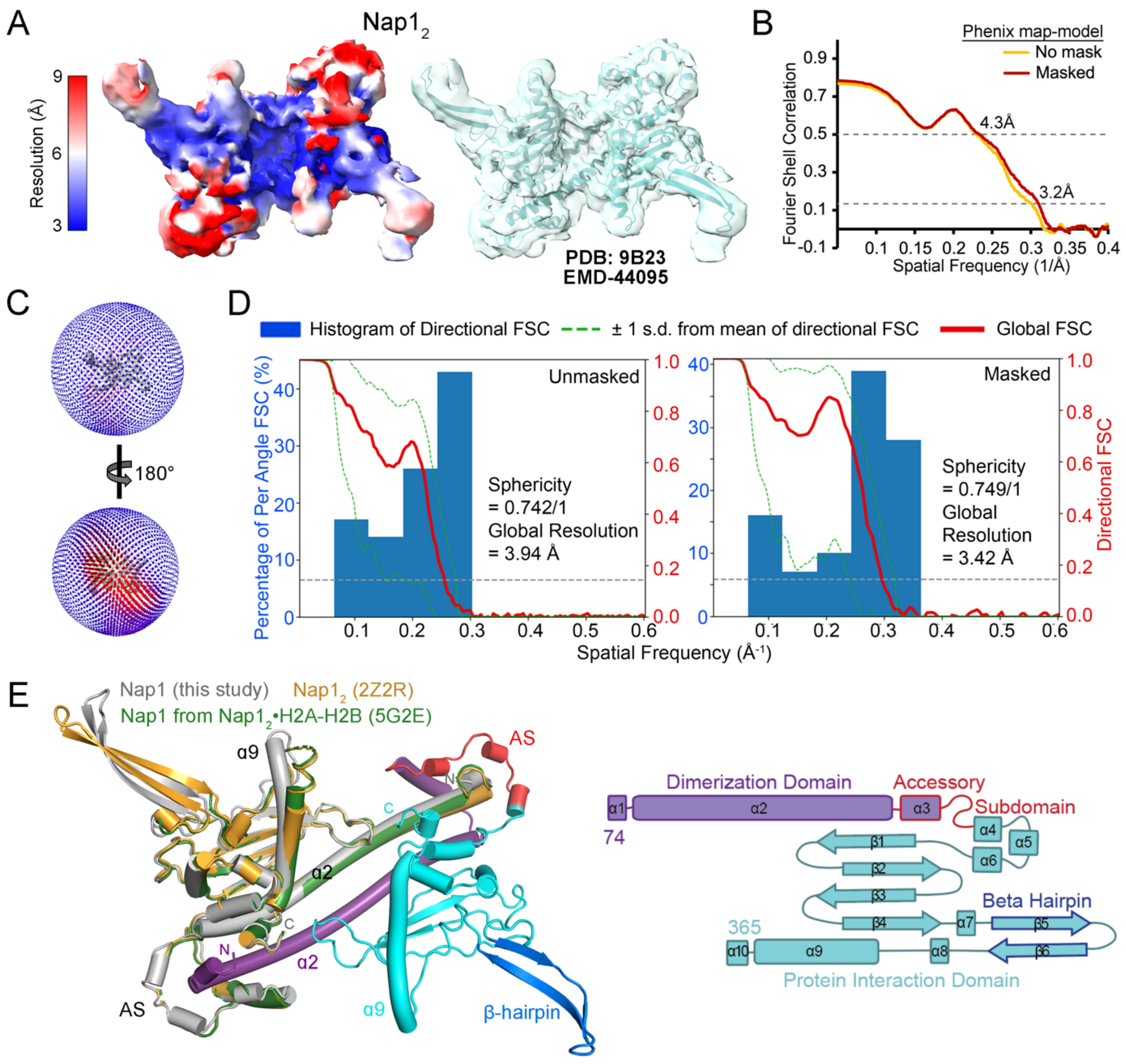


**Figure S4. Nap1$_2$: cryo-EM map details and structural comparison. (A)** The cryo-EM map of Nap1$_2$ (core; EMD-44095) colored by local resolution (left) and overlayed as transparent surface onto final model (right; PDB: 9B23). **(B)** Phenix map-to-model FSC curves for the main map. **(C)** 3D angular distribution of the particles that were used for reconstruction. The top orientation is same as in **(A)**. **(D)** Directional FSCs generated using the 3DFSC server (4), unmasked and masked by cryoSPARC refine mask. **(E)** The cryo-EM structure of Nap1$_2$ drawn with one monomer in gray, and the other subunit colored according to the schematic shown on the right. One subunit of the cryo-EM structure is aligned to one subunit of Nap1$_2$ from the crystal structures of Nap1$_2$•H2A-H2B (PDB: 5G2E (5); Nap1$_2$ is green and the H2A-H2B was not displayed) and of unliganded Nap1$_2$ (PDB: 2Z2R (6); gold). Root mean square deviation (r.m.s.d.) for the alignment with 5G2E is 1.1 Å (436 Cα atoms aligned) and with 2Z2R is 1.2 Å (454 Cα atoms aligned). The Nap1$_2$ regions with low local resolution in **(A)**, docked using AlphaFold-Multimer model, are also the regions that were unmodeled in the crystal structures; they include the accessory subdomain loop (AS, residues 170-180) and the N-terminal part of α9. There are small differences between the cryo-EM and crystal structures: the β-hairpins and the N-terminal parts of α9 are angled slightly differently and the α2 helix in the cryo-EM structure is slightly less bent.

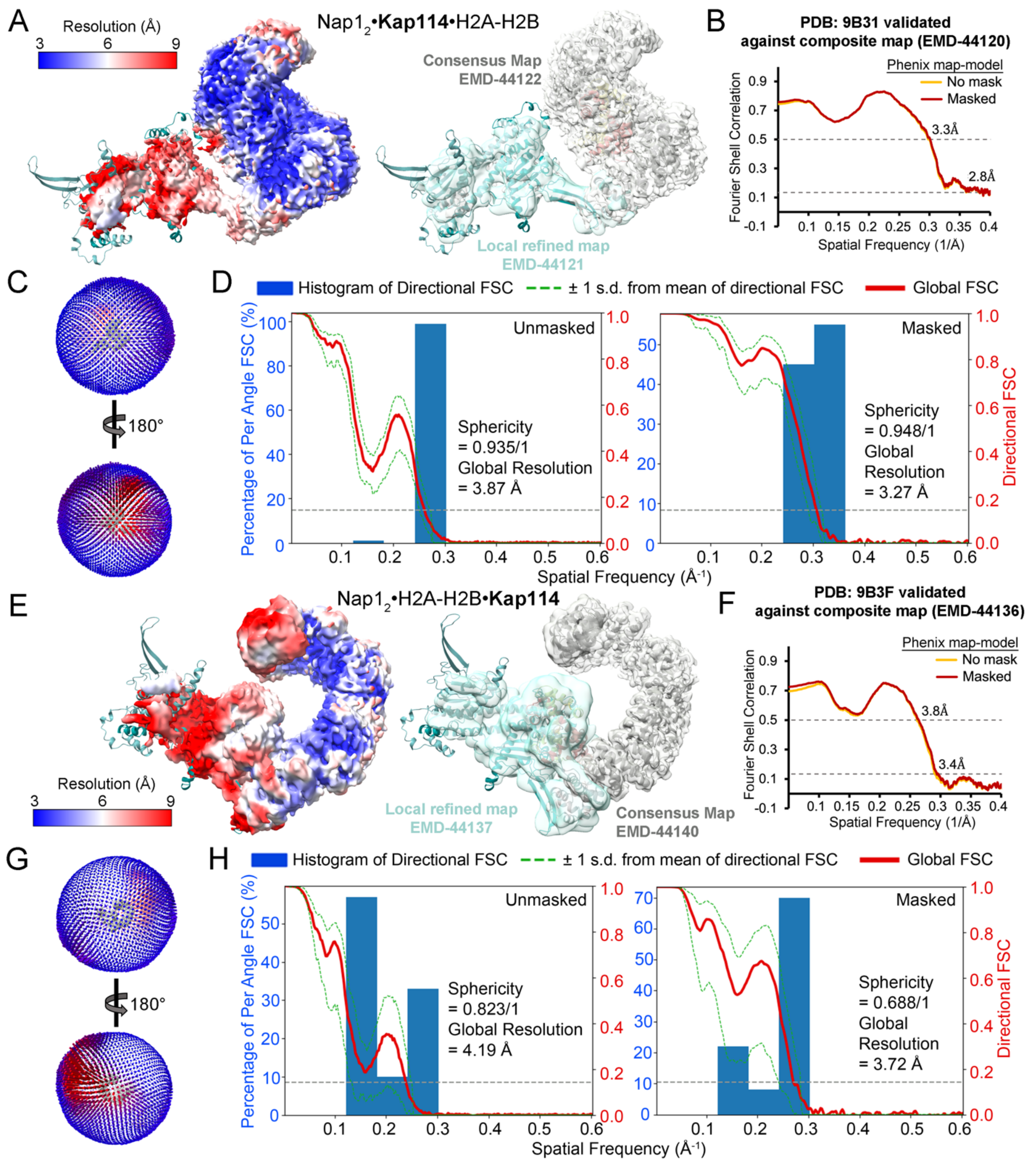


**Figure S5. Cryo-EM statistics and local resolution maps for the Kap114/Nap1$_2$/H2A-H2B structures. (A)** Left, the Nap1$_2$•**Kap114**•H2A-H2B consensus map (EMD-44122) colored by local resolution. Right, the same map (gray surface) and the locally refined map of the Nap1$_2$ region (blue; EMD-44121) are overlayed onto the final model (PDB: 9B31). **(B)** Phenix map-to-model FSC curves validated against the composite map (EMD-44120). **(C)** 3D angular distribution of the particles that were used for reconstruction. Top orientation is the same as in (**A**). **(D)** Directional FSCs generated using 3DFSC server (4), unmasked and masked by cryoSPARC refine mask. **(E-H)** As in **(A)-(D)** but for Nap1$_2$•H2A-H2B•**Kap114** (EMD-44140). The blue map in **(E)** is the locally refined map for Nap1$_2$•H2A-H2B (EMD-44137). The final model (PDB: 9B37) was validated against composite map (EMD-44136).

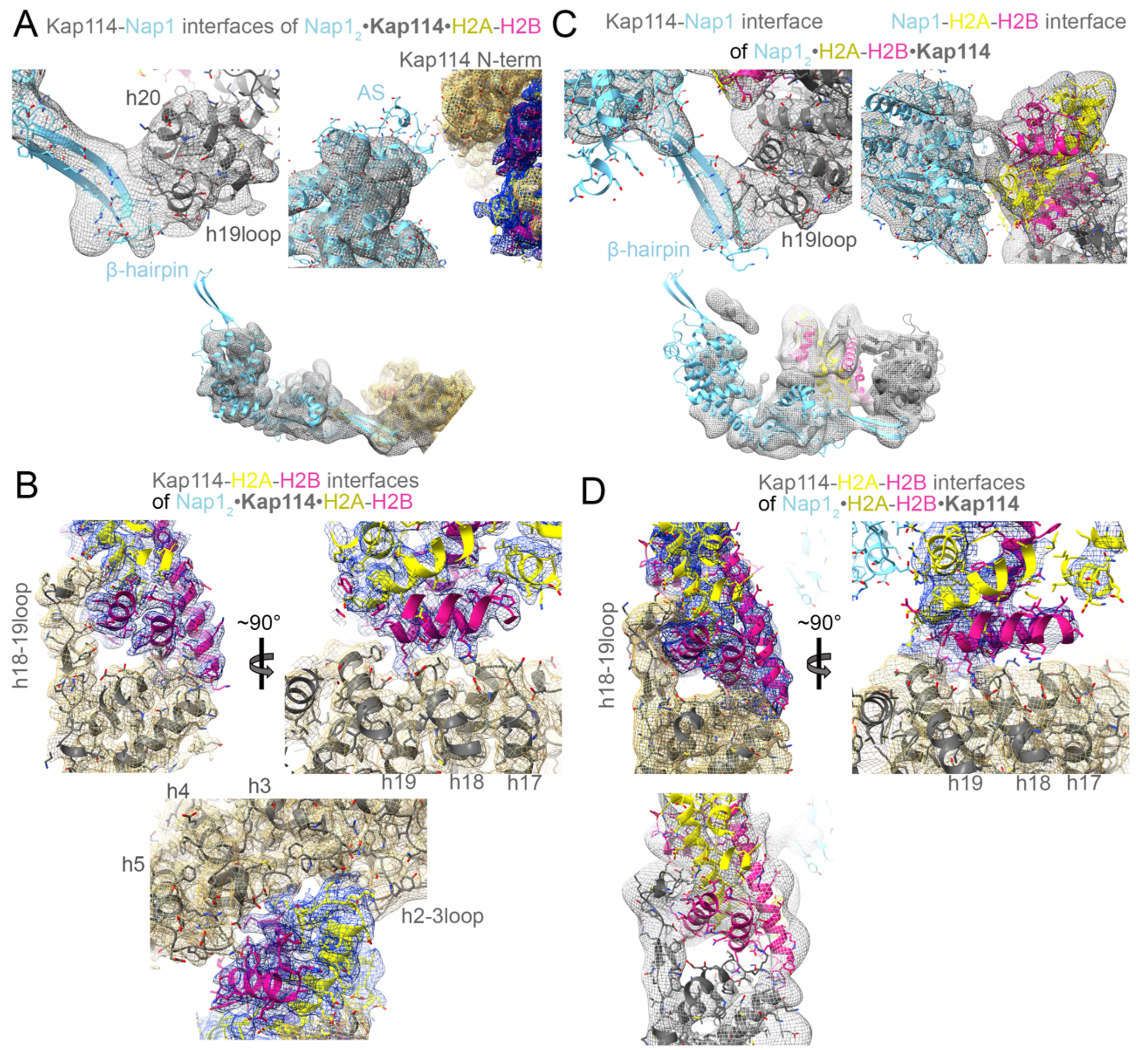


**Figure S6. Map quality at interfaces of the two Kap114/Nap1$_2$/H2A-H2B structures. (A)** The Nap1$_2$•**Kap114**•H2A-H2B structure (Nap1$_2$ (cyan), Kap114 (dark gray), H2A-H2B (yellow and pink)), shown in cartoon representation with sidechains in sticks, overlayed onto the local refined map (gray mesh) and the consensus map (Kap114, gold mesh and H2A-H2B, dark blue mesh). The top panel shows Kap114-Nap1$_2$ interfaces. The bottom panel shows the local refined map of the entire Nap1$_2$. **(B)** The Nap1$_2$•**Kap114**•H2A-H2B consensus map at the Kap114-H2A-H2B interfaces. **(C)** The Nap1$_2$•H2A-H2B•**Kap114** structure, oriented as in **(A)**, overlayed onto the local refined map for Nap1$_2$•H2A-H2B (gray mesh). Top, Kap114-Nap1$_2$ and Nap1$_2$-H2A-H2B interfaces. Bottom, map showing the entire Nap1$_2$ and H2A-H2B. **(D)** The Nap1$_2$•H2A-H2B•**Kap114** consensus map (Kap114, gold mesh and H2A-H2B, dark blue mesh) at the Kap114-H2A-H2B interfaces, orientated as in **(B)**. The local refined maps of both structures, shown in the bottom panels of **(A)** and **(C)**, suggest that the β-hairpins of the Nap1$_2$ subunit that does not contact Kap114 **(A)** or Kap114 and H2A-H2B **(C)** are very flexible and dynamic in both structures.

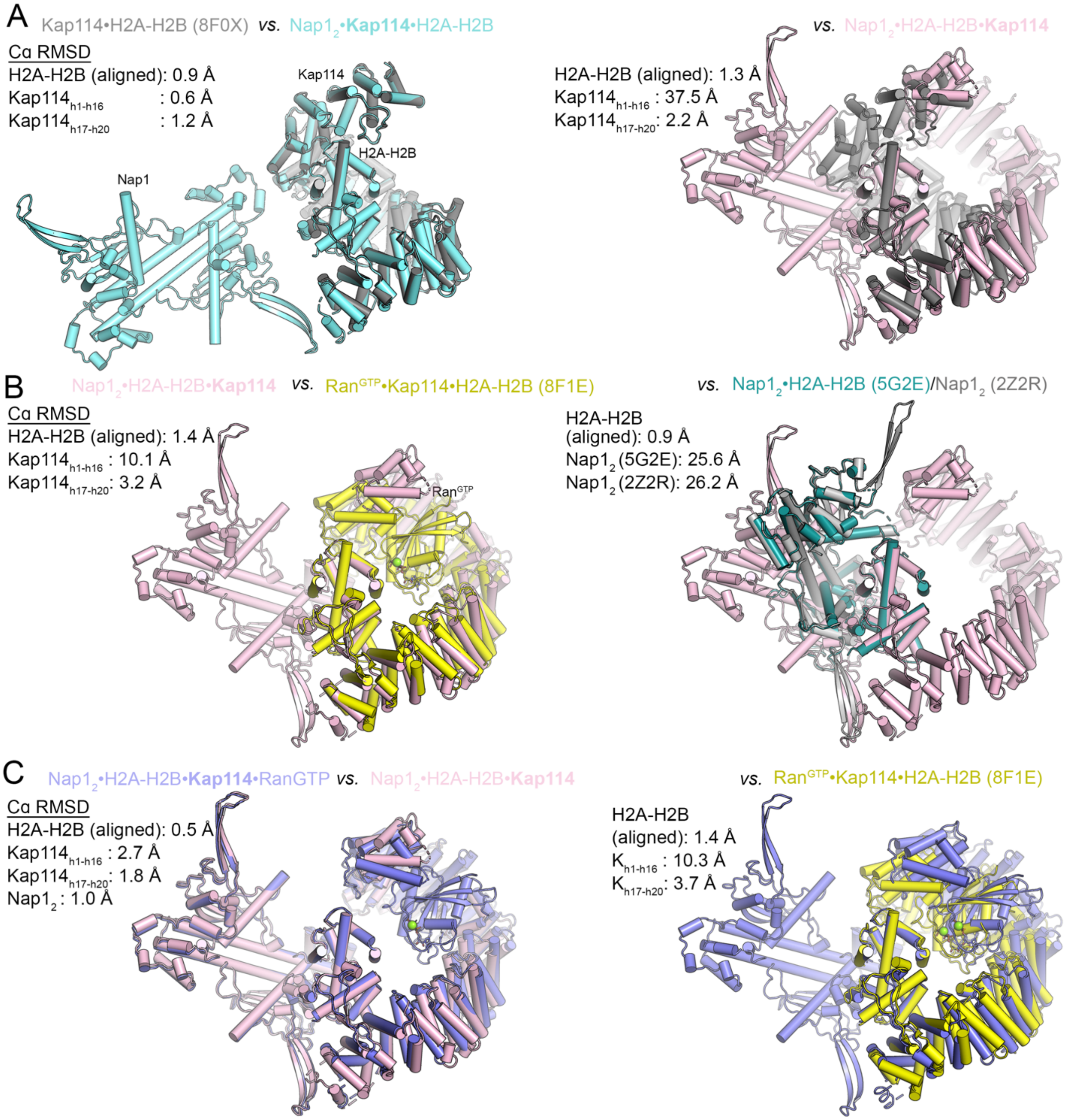


**Figure S7. Alignments of Kap114 and H2A-H2B from multiple structures.** The H2A-H2B heterodimers of various structures are aligned and the Cα r.m.s.d. values (calculated in PyMOL) of different molecules are reported. **(A)** Alignment of H2A-H2B heterodimers to superimpose Kap114•H2A-H2B (grey; PDB: 8F0X) with Nap1$_2$•**Kap114**•H2A-H2B (cyan; left) and Kap114•H2A-H2B with Nap1$_2$•H2A-H2B•**Kap114** (pink; right). **(B)** Alignment of H2A-H2B heterodimers to superimpose Nap1$_2$•H2A-H2B•**Kap114** (pink) with Ran$^{GTP}$•Kap114•H2A-H2B (yellow; PDB: 8F1E; left) and with Nap1$_2$•H2A-H2B (dark cyan; PDB: 5G2E; right). Unliganded Nap1$_2$ (PDB 2Z2R; grey) is overlayed onto Nap1$_2$•H2A-H2B to show putative locations of the unresolved β-hairpins in 5G2E (detailed analysis in Fig. S17). **(C)** Alignment of the H2A-H2B heterodimers of Nap1$_2$•H2A-H2B•**Kap114**•Ran$^{GTP}$ (purple) with Nap1$_2$•H2A-H2B•**Kap114** (pink; left) and with Ran$^{GTP}$•Kap114•H2A-H2B (yellow; right).

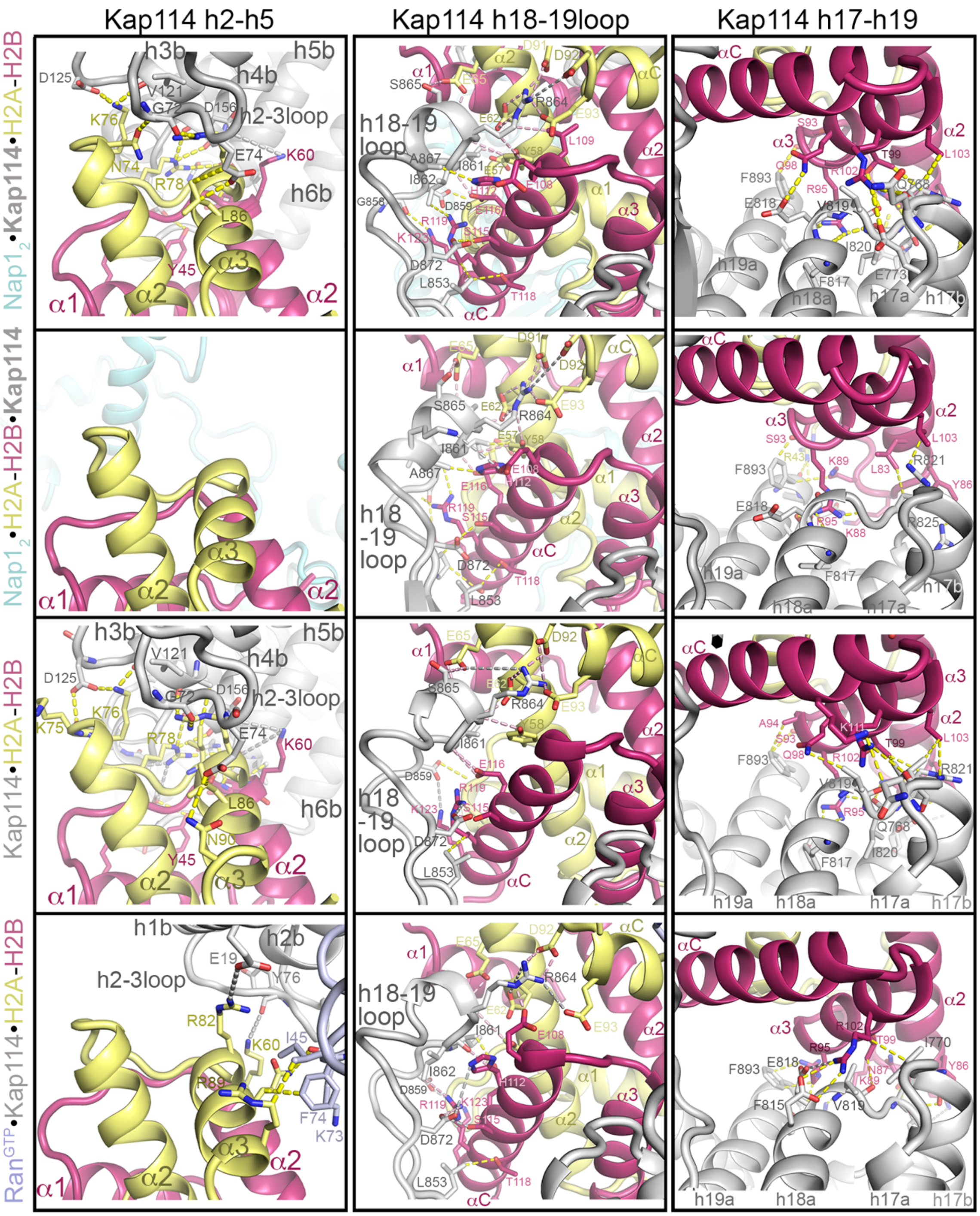


**Figure S8. Kap114-H2A-H2B binding interfaces in all available structures with Kap114 and H2A-H2B.** Top to bottom, Nap1$_2$•**Kap114**•H2A-H2B and Nap1$_2$•H2A-H2B•**Kap114** from this study and Kap114•H2A-H2B (PDB: 8F0X) and Ran$^{GTP}$•Kap114•H2A-H2B (PDB: 8F1E) (7). Left to right, interactions of H2A-H2B with HEAT repeats 2-5 of Kap114, or lack thereof, and the persistent interactions with h18-19 loop of Kap114 and h17-h19.

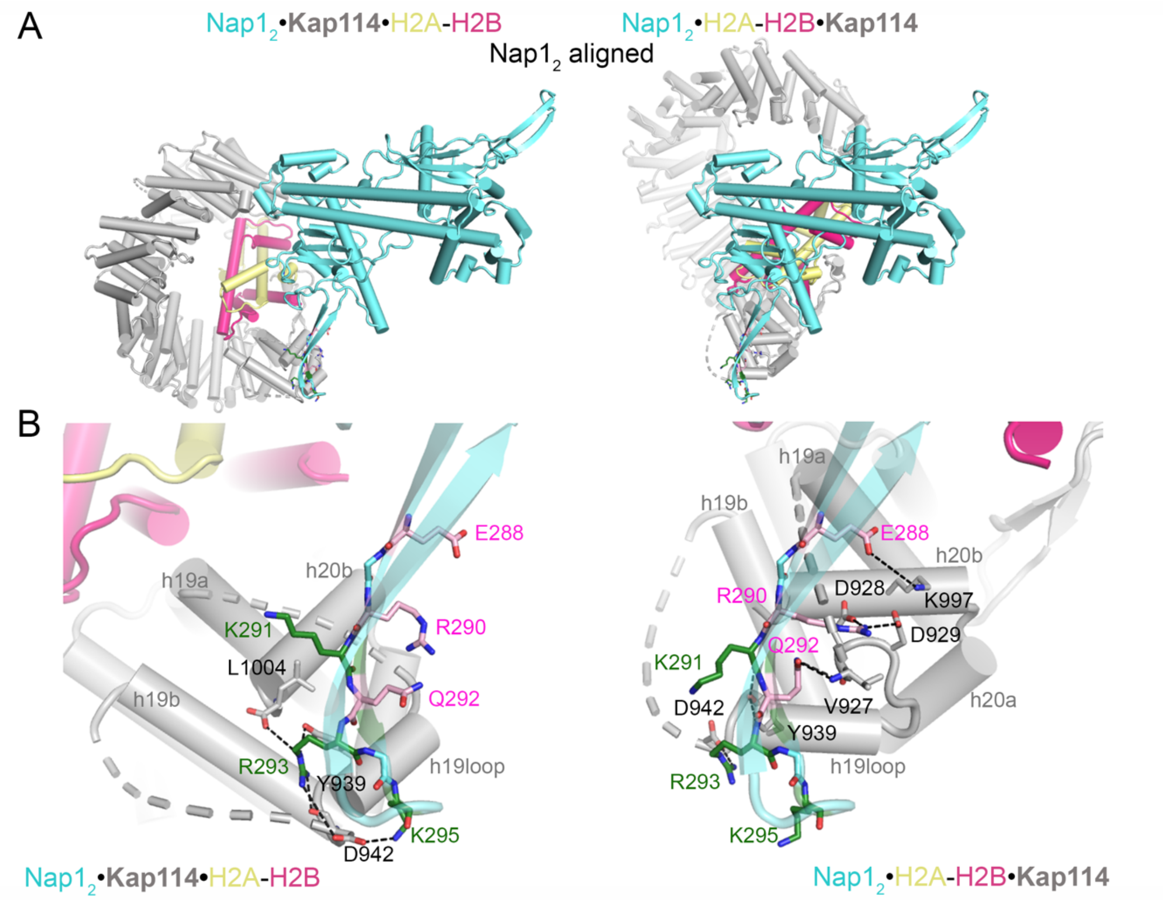


**Figure S9. Kap114 contacts two different faces of the Nap1$_2$ β5 strand in Nap1$_2$•*Kap114*•H2A-H2B *vs.* Nap1$_2$•H2A-H2B•*Kap114*. (A)** The two Kap114/Nap1$_2$/H2A-H2B (gray/cyan/yellow-pink) structures aligned by Nap1$_2$. **(B)** Zoom-in views of the Kap114-Nap1$_2$ interfaces of the two structures, oriented as in **(A)**. Kap114-Nap1 contacts are shown as black dashed lines. In the left panel, the Kap114 in Nap1$_2$•**Kap114**•H2A-H2B contacts residues K291, R293, K295 (green sticks) on one side of the β5 strand of one subunit of Nap1$_2$. These residues were mutated in the Nap1 $\beta_{KRK}$ mutant that was used in **Figure 3C**. In the right panel, the Kap114 in Nap1$_2$•H2A-H2B•**Kap114** contacts residues E288, R290, Q292 (pink sticks) on a different the side of the Nap1 β5 strand. These residues were mutated in the Nap1 $\beta_{ERQ}$ mutant that was used in **Figure 3C**.

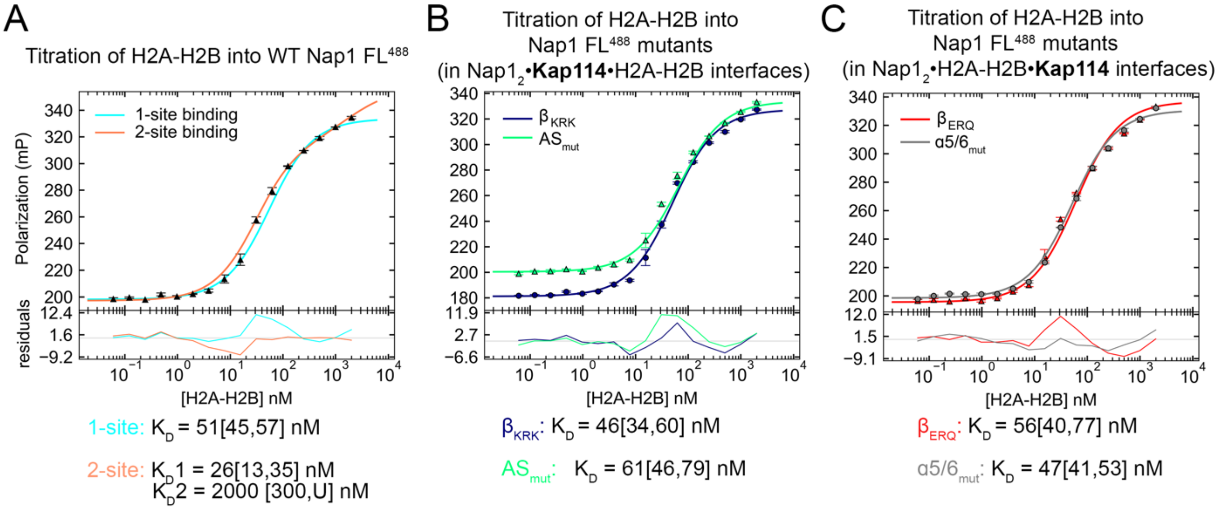


**Figure S10. Affinity of Nap1 WT and mutants for H2A-H2B. (A)** FP titration of H2A-H2B into 10 nM $Nap1_2$ FL labeled with XFD488 (Nap1 $FL^{488}$). Data points are averages ± s.d. of triplicate measurement. The lines show data fitted with 1-site or 2-site binding and residuals are plotted below. Dissociation constants are recorded with the 95% confidence interval obtained by error-surface projection method in brackets. The data is better fitted with 2-site binding, which is consistent with previous work that reported human $Nap1_2$ binding two copies of H2A-H2B, one bound to the C-terminal acidic tails and one to the core (8, 9). **(B)** FP experiment like **(A)** but with Nap1 proteins mutated at the Kap114-$Nap1_2$ interface of the $Nap1_2$•**Kap114**•H2A-H2B structure. The Nap1 mutants are the $\beta_{KRK}$ β-hairpin mutant and the $AS_{mut}$ mutant. A 1-site binding model was used as data could not be fitted confidently with 2-site binding. **(C)** FP experiment like in **(B)** but with Nap1 mutated at the Kap114-Nap1 or Nap1-H2A-H2B interfaces of the $Nap1_2$•H2A-H2B•**Kap114** structure. The Nap1 mutants are the $\beta_{ERQ}$ β-hairpin mutant and the $\alpha5/6_{mut}$ mutant. A 1-site binding model was used for fitting as data could not be fitted confidently with 2-site binding. All Nap1 mutants bound H2A-H2B with high affinity in the low nM range.

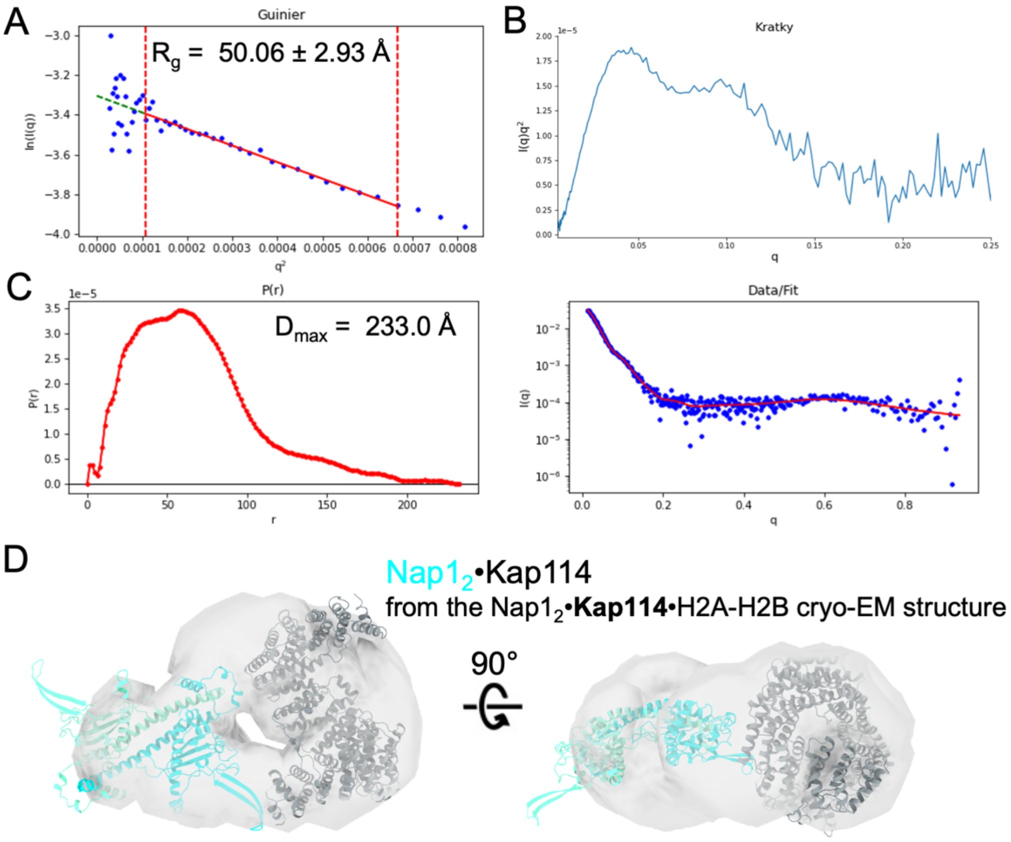


**Figure S11. Small Angle X-Ray Scattering (SAXS) analysis of the binary Nap1$_2$•Kap114 complex. (A)** The linear region of Guinier plot with calculated radius of gyration ($R_g$) fit value in Å. Blue dots represent the data while the red dashed lines are the boundaries of the fitted red line. **(B)** Kratky plot of the Nap1$_2$•Kap114 complex suggests that this complex is a well-folded globular protein. **(C)** Left, the pair distribution function, P(r), along with the maximum particle size ($D_{max}$) determined from the maximum pair distance on the plot. Right, the scattering profile in blue, and the fit generated during the pair distance function in red. **(D)** Shown in two orientations, the light gray *ab-initio* electron density (reconstructed using DENSS) of the Nap1$_2$•Kap114 SAXS sample, with Nap1$_2$•Kap114 (cyan/gray) from the Nap1$_2$•**Kap114**•H2A-H2B cryo-EM structure docked into the density. The reconstructed SAXS envelope of Kap114•Nap1$_2$ matches well with the Kap114-Nap1$_2$ arrangement in the ternary Nap1$_2$•**Kap114**•H2A-H2B structure. This is consistent with mutagenesis/pull-down experiments in Figure 3C and D showing the importance of Nap1$_2$ β-hairpin binding to the Kap114 h19loop even when H2A-H2B is not present, suggesting that the same Kap114-Nap1 binding mode in the Nap1$_2$•Kap114•H2A-H2B structure is also used to form the binary Kap114•Nap1 complex.

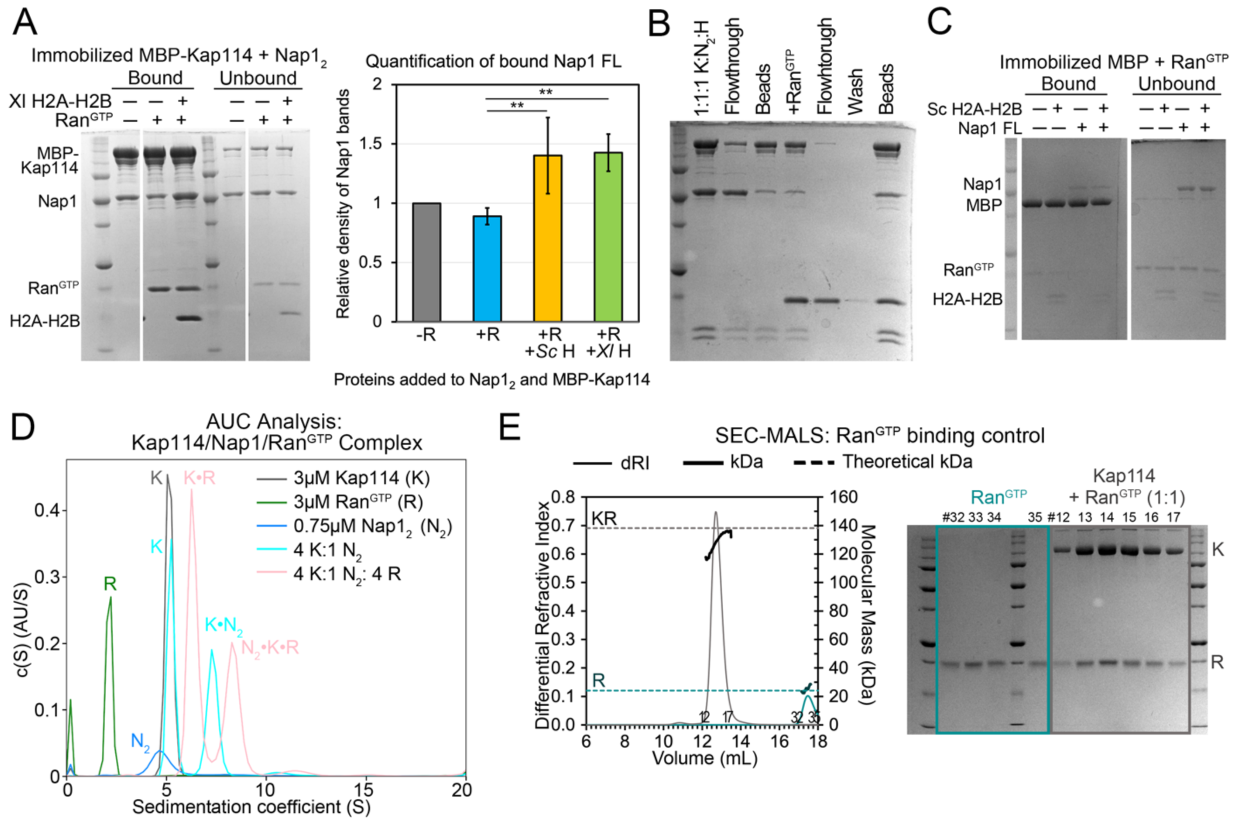


**Figure S12. Interactions between Kap114, Nap$1_2$ and H2A-H2B in the presence of Ran$^{GTP}$. (A)** Pull-down assay using 1:1:1:1 molar ratio of immobilized MBP-Kap114 (1 μM), Nap$1_2$ FL ± *Xl* H2A-H2B ± Ran$^{GTP}$. Left, bound/unbound proteins visualized by Coomassie-stained SDS-PAGE. Right, quantification of the Kap114-bound Nap1 band intensities from triplicate experiments with *Sc* (gel in **Figure 4B**) or *Xl* H2A-H2B (H) ± Ran$^{GTP}$ (R). ** indicate p-value <0.01. MBP-Kap114 pulled down more Nap1 when both H2A-H2B and Ran$^{GTP}$ are present. These results are consistent with SEC-MALS data showing a smaller shoulder at ~ 12.2 mL of the Nap$1_2$•H2A-H2B•**Kap114**•Ran$^{GTP}$ peak (green) than in the Nap$1_2$•H2A-H2B•**Kap114** peak (orange) in **Figure 4C**. **(B)** The same pull-down assay as in **Figure 4B**, except 3 μM Ran$^{GTP}$ was added after pre-assembly of the MBP-Kap114/Nap$1_2$/H2A-H2B (K:$N_2$:H) complex and immobilization on beads. Individual steps of the binding assay are visualized by SDS-PAGE. The quaternary Kap114/Nap$1_2$/H2A-H2B/Ran$^{GTP}$ complex formed regardless of the order of protein addition, and no Nap1 or H2A-H2B was dissociated by Ran$^{GTP}$. **(C)** Control pull-down assays of 1 μM MBP (immobilized) with equimolar H2A-H2B and/or Nap1 FL in the presence of Ran$^{GTP}$. **(D)** AUC analysis. Plots of the c(s) distributions of unliganded proteins, Kap114 (K, 5.2 S), Nap$1_2$ core ($N_2$, 4.9 S) and Ran$^{GTP}$ (R, 2.1 S), and mixtures of the proteins with indicated molar ratios. The species corresponding to the individual peaks are labeled. Ran$^{GTP}$ binding increased sedimentation coefficient similarly for unliganded Kap114 (K, 5.2 → K•R, 6.3S) and for Kap114•Nap$1_2$ (K•$N_2$, 7.3 → $N_2$•K•R, 8.3S). The Kap114•Ran$^{GTP}$ and Nap$1_2$•Kap114•Ran$^{GTP}$ complexes had estimated molecular weights of 138 kDa and 206 kDa, respectively, consistent with equimolar complexes. **(E)** SEC-MALS analysis, the Ran$^{GTP}$ binding control for the experiment in **Figure 4C**. The curves drawn with thin lines are differential refractive index, those with thick lines are molecular mass and the dashed lines indicate theoretical molecular mass of the predicted complexes. Fractions from the two peaks (indicated by the respective colored boxes) were visualized by SDS-PAGE and Coomassie staining.

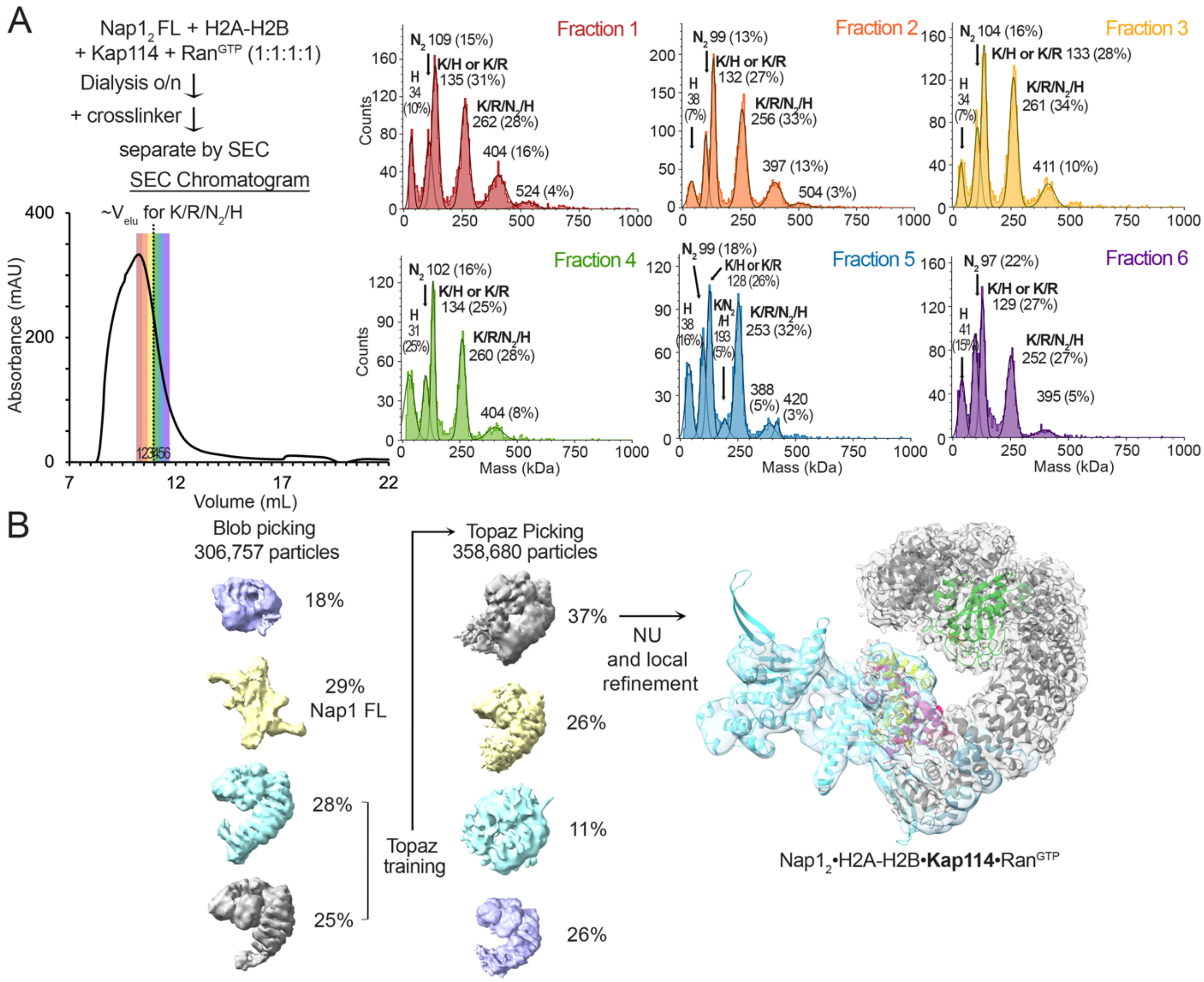


**Figure S13. Cryo-EM sample of the Kap114/Nap1$_2$/H2A-H2B/Ran$^{GTP}$ quaternary complex. (A)** Top left, schematic of the assembly/crosslinking of the Kap114 (K), Nap1 FL (N$_2$), H2A-H2B (H) and Ran$^{GTP}$ (R) mixture and the subsequent size-exclusion chromatography (SEC). Bottom left, SEC chromatogram with fractions 1-6 colored red to purple. The typical elution volume of a non-crosslinked complex that contains all four proteins is marked with a dotted line. The six SEC fractions were subjected to mass photometry; traces are shown on right. The six panels show mass photometry data with mean masses (kDa) and relative populations (%) indicated above the fitted gaussian peaks, along with the likely protein or complex that correspond to the approximate masses. Fraction 3 is most enriched with the complex containing all four proteins and was used or cryo-EM grid preparation. The ~380-410 kDa species may be crosslinked complexes of K/R/N$_2$/H with another copy of K:R. Without crosslinking, no such large assembly was observed, by AUC or SEC-MALS. **(B)** Particle distribution of Cryo-EM data obtained for the Kap114/Nap1$_2$/H2A-H2B/Ran$^{GTP}$ quaternary complex. Blob picking was used first and then the particles that contain Kap114 and Ran$^{GTP}$ were used for Topaz training. Topaz-picked particles were cleaned up and submitted to 3D reconstruction to obtain 4 maps. The population with density (albeit poor) for H2A-H2B and Nap1$_2$ was used for non-uniform (NU; gray map) and local refinement (cyan map) to obtain the final maps. The maps were overlayed onto the final Nap1$_2$•H2A-H2B•Kap114•Ran$^{GTP}$ structure. Nap1$_2$ is cyan, H2A yellow, H2B red, Kap114 gray and Ran is green.

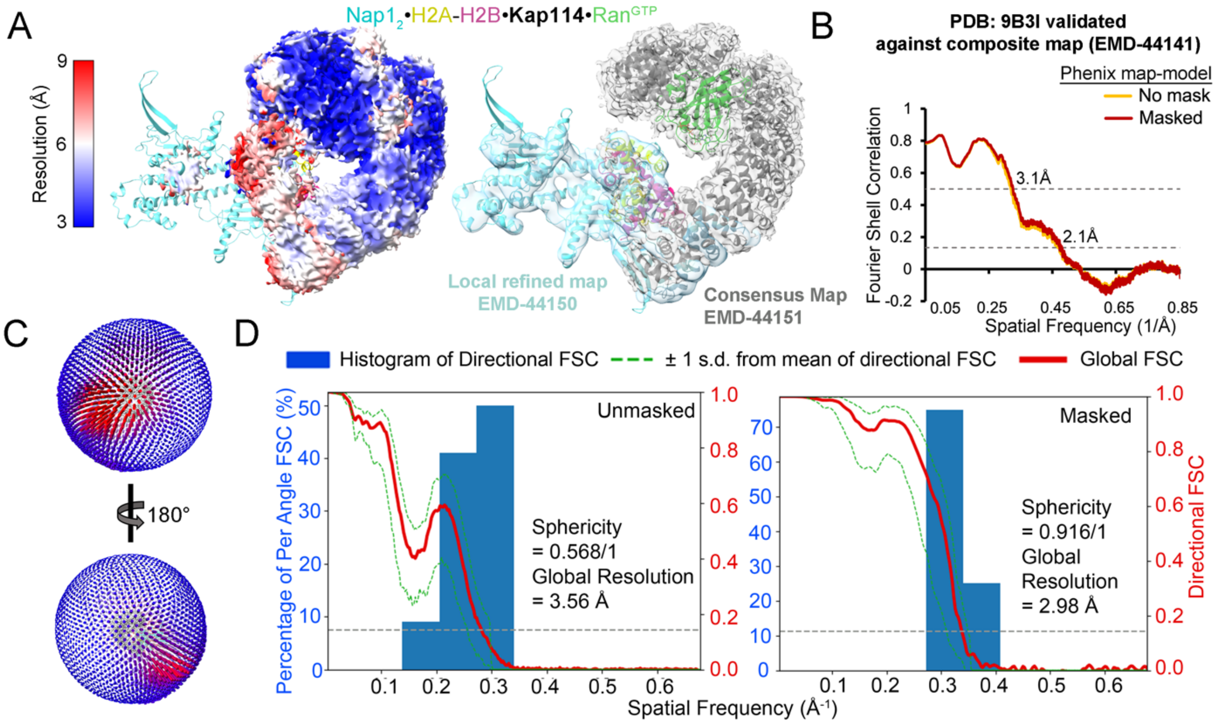


**Figure S14. Details of the Nap1$_2$•H2A-H2B•*Kap114*•Ran$^{GTP}$ cryo-EM map. (A)** Left, the final map from NU refinement (EMD-44151) colored by local resolution. Right, the consensus map shown as gray transparent surface and the local refined map for Nap1$_2$ and H2A-H2B shown as cyan transparent surface (EMD-44150), overlayed onto the final structure (PDB: 9B3I). **(B)** Phenix map-to-model FSC curves for the composite map (EMD-44141). **(C)** 3D angular distribution of the particles used for reconstruction. Top orientation is the same as in **(A)**. **(D)** Directional FSCs generated using 3DFSC server (4), unmasked, and masked by cryoSPARC refine mask for the NU refined map.

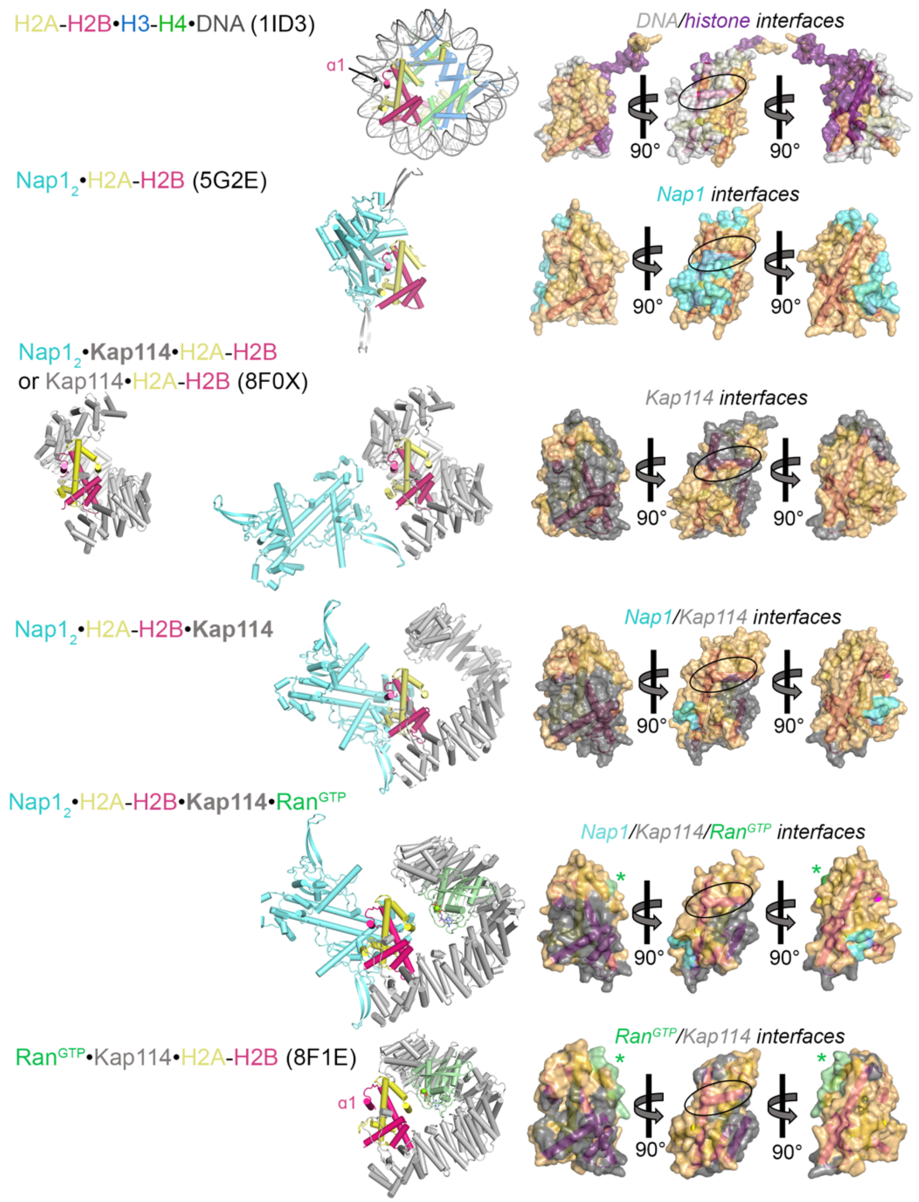


**Figure S15. H2A-H2B interfaces.** Left panel, top to bottom: Structures of the nucleosome (1ID3), Nap1$_2$•H2A-H2B (5G2E), Nap1$_2$•**Kap114**•H2A-H2B and Kap114•H2A-H2B (8F0X), Nap1$_2$•H2A-H2B•**Kap114**, Nap1$_2$•H2A-H2B•**Kap114**•Ran$^{GTP}$ and Ran$^{GTP}$•Kap114•H2A-H2B (8F1E). Right panel, top to bottom: three views of the semi-transparent H2A-H2B surface with cartoon representation underneath, for the corresponding structures in the left panel. Binding interfaces (PDBePISA) are colored according to binding partners. The green asterisks (*) indicate transient Ran$^{GTP}$-H2A-H2B contacts. The $\alpha$1 helix of H2B, labeled or indicated by ovals in one view of the H2A-H2B surfaces, is one of the nucleosomal DNA-binding regions. H2B $\alpha$1 is buried by Nap1$_2$ in Nap1$_2$•H2A-H2B, and by Kap114 in Kap114•H2A-H2B and Nap1$_2$•**Kap114**•H2A-H2B. In Nap1$_2$•H2A-H2B•**Kap114** and Nap1$_2$•H2A-H2B•**Kap114**•Ran$^{GTP}$, Nap1$_2$ does not make <4 Å contacts with H2B $\alpha$1 but is nearby (< 10 Å away), limiting access of other macromolecules to the histone. In contrast, H2B $\alpha$1 is exposed in Ran$^{GTP}$•Kap114•H2A-H2B, likely explaining why histone is not effectively chaperoned in this complex. The proximity of the bound Nap1$_2$ to H2B $\alpha$1 in the quaternary Nap1$_2$•H2A-H2B•**Kap114**•Ran$^{GTP}$ complex likely contributes to successful chaperoning of H2A-H2B in the presence of Ran$^{GTP}$.

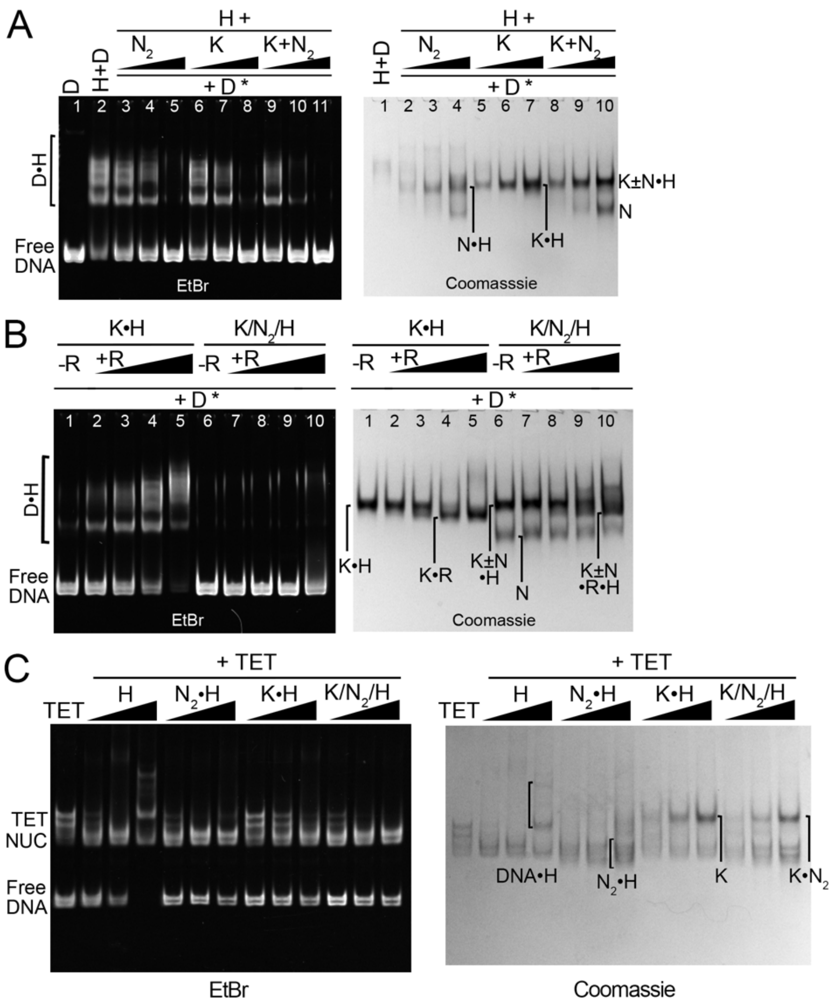


**Figure S16. DNA competition and nucleosome assembly assays with Kap114 and Nap1**. **(A)** DNA competition assay with pre-assembled complexes of 4 µM H2A-H2B (H) first added to 1, 2 or 4 µM of Nap1$_2$ FL (N$_2$). Kap114 (K) or K+N$_2$ before addition of 1 µM DNA at the end. Samples were visualized by Native-PAGE and EtBr (left) and Coomassie (right) staining. **(B)** As in **A**, DNA competition assay with pre-assembled complexes of 4 µM H added to 4 µM of N$_2$, K or K+N$_2$ and 1, 2, 4 or 8 µM RanGTP (R), which then 1 µM DNA was added at the end. **(C)** 0.5, 1 and 2 µM H was pre-mixed with equimolar N$_2$ and/or K before adding to 1 µM tetrasomes (assuming 100% formation of DNA•(H3-H4)$_2$). The EtBr gel shows that H2A-H2B alone can incorporate into tetrasomes (TET) to form nucleosomes (NUC)[ but higher-order aggregates formed in the presence of excess H2A-H2B. Nap1$_2$•H2A-H2B deposited histones to form nucleosomes without forming aggregates. Kap114•H2A-H2B inhibited H2A-H2B deposition but the presence of Nap1$_2$ in the Kap114/Nap1$_2$/H2A-H2B complex reversed the inhibition by Kap114, allowing efficient histone deposition to form nucleosomes. The Kap114 inhibition of H2A-H2B deposition was less than previously reported (10), probably due to *Sc* H2A-H2B used here instead of the *Xl* H2A-H2B used in (4).

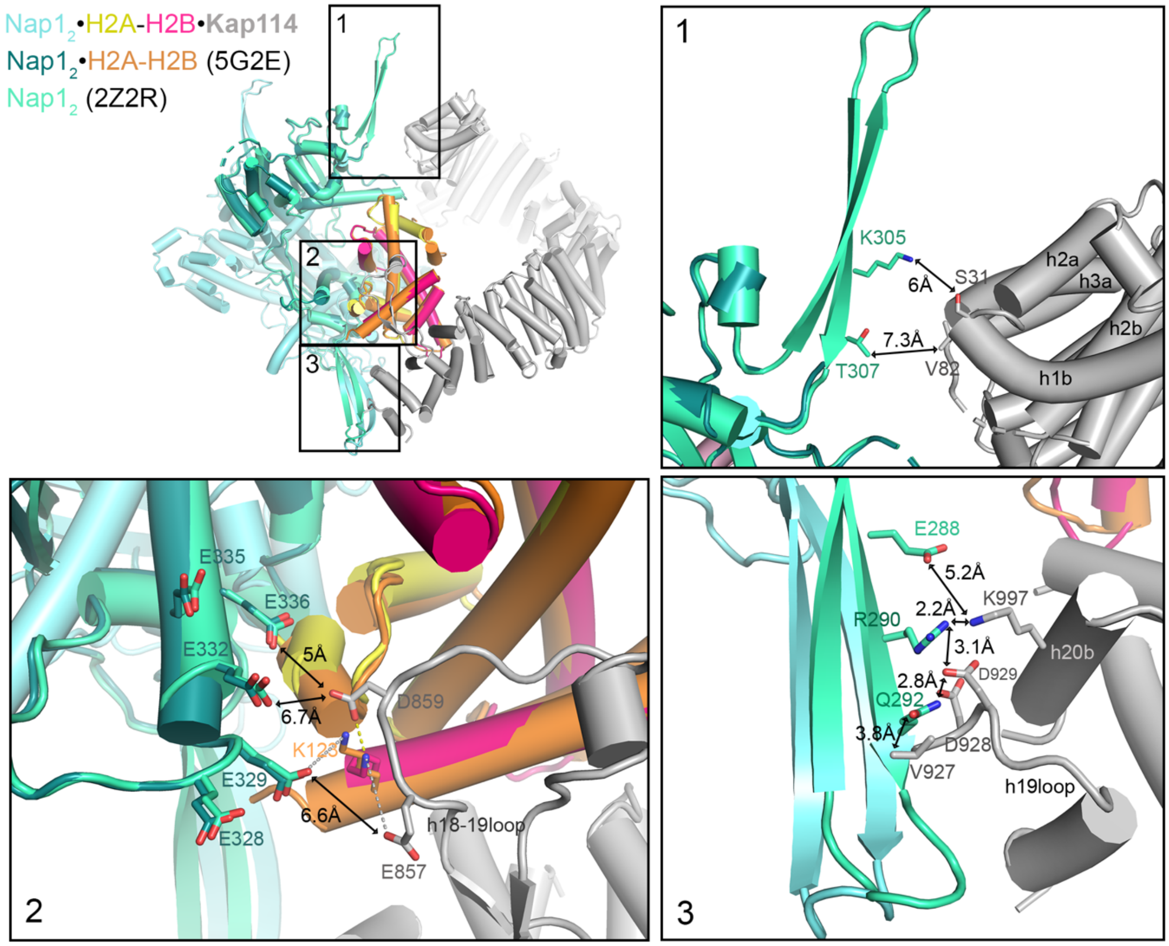


**Figure S17. Analysis of the superimposed Nap1$_2$•H2A-H2B and Nap1$_2$•H2A-H2B•*Kap114* structures.** Top left, the overall view of the superimposed Nap1$_2$•H2A-H2B and Nap1$_2$•H2A-H2B•**Kap114** (H2A-H2B aligned**,** as in the right panel of Fig. S7B). Here, Nap1$_2$•H2A-H2B•**Kap114** is in cyan•yellow-pink•gray, Nap1$_2$•H2A-H2B (5G2E) is in dark teal•orange. Nap1$_2$ (2Z2R; green) is superimposed onto 5G2E to show locations of the disordered β-hairpins of the latter structure. The numbered boxes refer to three separate regions where details are shown in the other panels. The top right panel shows region 1 where the β-hairpin of one Nap1$_2$ subunit is ~7 Å away from the h1-2loop and h2-3loop of Kap114. The bottom left panel shows region 2, which is at the Nap1$_2$-histone interfaces of the superimposed Nap1$_2$•H2A-H2B and Nap1$_2$•H2A-H2B•**Kap114** structures. Nap1$_2$ of the superimposed Nap1$_2$•H2A-H2B does not clash with the Kap114 h18-h19loop, which makes many interactions with the Nap1$_2$-bound H2A-H2B of Nap1$_2$•H2A-H2B•**Kap114**. However, superposition unfavorably places many acidic residues of Nap1$_2$•H2A-H2B close to acidic chains of Kap114 h18-h19loop. The bottom right panel shows region 3 where the other Nap1$_2$ β-hairpin of the superimposed Nap1$_2$ (2Z2R)/Nap1$_2$•H2A-H2B (5G2E) is very close to the Kap114 h19loop and helix h20b of Nap1$_2$•H2A-H2B•**Kap114**. We show the Nap1$_2$ β-hairpin side chains of E288, R290 and Q292; some of them may contact Kap114 while others may clash with the Kap114 h19loop conformation of Nap1$_2$•H2A-H2B•**Kap114**. The proximity of similar negative charges in region 2 and potential clashes in region 3 may suggest that the superimposed model is unfavorable, possibly explaining its absence in our cryo-EM analysis. Reorientation of Nap1$_2$ and H2A-H2B to what we see in Nap1$_2$•H2A-H2B•Kap114 may be necessary for stable interactions of the three proteins.

**Protein constructs, expression, and purification**

Kap114 was cloned into vectors pGEX-4T3 (Cytiva) and pMalE (New England BioLabs). pGEX-4T3 was modified with a TEV cleavage site between the GST tag and Kap114 whereas pMalE was modified with a $His_6$-tag immediately N-terminus of MBP and a TEV cleavage site after the MBP. Mutant proteins were generated by site-directed mutagenesis or blunt-end ligations.

GST-Kap114 proteins were expressed and purified as previously described (10). MBP-Kap114 was expressed in BL21 Gold cells grown in LB media and protein expression was induced with 0.5 mM IPTG for 17 hr at 18 ºC. Cells were harvested by centrifugation at 4000 g (Sorvall BP6) and resuspended in lysis buffer containing 50 mM HEPES 7.0, 150 mM NaCl, 10% (v/v) glycerol, 5mM DTT, 1 mM benzamidine, 10 µg/mL leupeptin and 50 µg/mL AEBSF and frozen. Thawed bacteria cells were lyzed using Emulsiflex homogenizer, the lysate clarified by centrifugation at 20000 rpm for 40 min at 4 °C (Sorvall RC6) and supernatant added to amylose beads (New England Biolabs). The beads were briefly washed with lysis buffer with NaCl added to 300 mM. MBP-Kap114 was then eluted with buffer containing 50 mM HEPES 7.0, 50 mM NaCl, 10% (v/v) glycerol, 20 mM maltose and 2 mM DTT and further purified by ion exchange using HiTrap Q HP column (Cytiva) in 25 mM Bis-Tris pH 6.5, 0 to 1M NaCl, 10% (v/v) glycerol, 2 mM DTT. MBP-Kap114 was subjected to a last purification step over SEC using a HiLoad Superdex 200 column (Cytiva) in Assay Buffer containing 20 mM HEPES pH 7.4, 150 mM NaCl, 2 mM $MgCl_2$, 10% (v/v) glycerol and 2 mM DTT.

Nap1 FL (C200A, C249A, C272A mutation for specific labeling on 414C) cloned into a pHAT4 vector was a gift from Sheena D'Arcy. Nap1 mutants were generated by site-directed mutagenesis using Phusion polymerase (Thermo Fisher) or CloneAmp HiFi PCR premix (Takara Bio). Nap1 FL (WT and mutants) were expressed in BL21 gold cells grown in 2X YT media and protein expression was induced by 0.5 mM IPTG for 17 hr at 18 ºC. Cells were harvested by centrifugation and resuspended in Nap1 lysis buffer containing 20 mM Tris pH 7.5, 1 M NaCl,

15% (v/v) glycerol, 1 mM DTT, 1 mM benzamidine, 10 µg/mL leupeptin and 50 µg/mL AEBSF. Thawed cells were lysed, and the clarified lysate was supplemented with 5 mM imidazole pH 7.8 and added to Ni-NTA agarose (Qiagen), which were washed with the Nap1 lysis buffer with 5 mM imidazole, pH 7.8. The beads were further washed with buffer containing 20 mM HEPES pH 7.4, 300 mM NaCl, 15% (v/v) glycerol, 1 mM 2-mercaptoethanol with 25 mM imidazole, pH 7.8. Bound protein was eluted with the same buffer supplemented to 250 mM imidazole and concentrated to ~10 mL. 1 mg of TEV protease was then added to the concentrated protein, overnight, at 4 ºC. The protein mixture was diluted and passed through Ni-NTA beads to remove TEV and the His tag. Nap1 is further purified using HiTrap Q and Superdex S200.

Nap1 core (residues 75-365), cloned into a pET15b vector, was a gift from Karolin Luger. His-Nap1 core proteins were expressed and purified as previously described (11). For pull-down assays, thrombin cleavage was performed overnight at 4 ºC to remove the His-tag and untagged protein purified by SEC (Superdex S200 increase column (Cytiva)) in Assay Buffer. Lyophilized *Sc* and *Xl* histones were obtained from The Histone Source and refolded according to established protocol (12). All assays in this paper is performed with *Sc* H2A-H2B unless otherwise stated (Figure S2A and S12A).

*Sc* $Ran^{GTP}$ (Gsp1 residues 1-179, with Q71L mutation to stabilize the GTP bound state) is expressed and purified as described previously with the addition of a TEV cleavage step (13). Briefly, Ran was expressed in BL21 gold cells, induced with 0.5 mM IPTG for 12 hr at 20 °C. Cells were resuspended in buffer containing 50 mM HEPES pH 7.4, 2 mM MgOAc, 200 mM NaCl, 10 % (v/v) glycerol, 5 mM imidazole pH 7.8, 2 mM 2-mercaptoethanol, 1 mM benzamidine, 10 µg/mL leupeptin, 50 µg/mL AEBSF. Thawed cells were lysed, and the clarified lysate was incubated with Ni-NTA beads, which were washed. Ran was eluted with buffer containing 50 mM HEPES pH 7.4, 2 mM MgOAc, 50 mM NaCl, 10% (v/v) glycerol, 250 mM imidazole pH 7.8 and 2 mM 2-mercaptoethanol, concentrated, and treated with 1 mg of TEV overnight incubation at 4 °C. TEV-cleaved $Ran^{GTP}$ was purified by ion-exchange using HiTrap SP HP column (Cytiva) in buffer containing 20 mM HEPES pH 7.4, 0 to 1 M NaCl, 4 mM MgOAc, 10% (v/v) glycerol and clean protein was flash-frozen and stored in -80 °C. $Ran^{GTP}$ activity was verified through binding assays with various importins that GTP is bound in the purified Ran and no additional GTP loading steps were needed.

**Mass photometry for cryo-EM samples**

All protein fractions were analyzed using a Refeyn TwoMP Mass Photometer. The laser was warmed up for an hour before use. During that waiting period, the glass slide was washed by alternating between Milli-Q filtered water, isopropanol, and then Milli-Q water, twice. A 2 by 4 strip of wells were adhered to the middle of the glass slide. Immersion oil was applied onto the lens before placing the glass slide on top. Prior to the measurement, 2 mg/mL BSA was diluted 100-fold with freshly filtered buffer from the buffer used in S200 of the cryo-EM complex. BSA was used for generating the calibration curve, using an additional 10-fold dilution: we first added 16.2 µL of buffer into a well and brought it to focus then mixed in 1.8 µL of BSA and 60 s movies were recorded. Fractions were diluted to under 7000 counts for collection in a similar manner as the BSA. Data was processed through Gaussian fitting to provide the peak mass and relative populations using the Refeyn analysis software.

**Cryo-EM data collection**

A 48-hour data collection with the best grid of the Kap114/H2A-H2B/Nap1 core complex was performed at the UT Southwestern Cryo-Electron Microscopy Facility (CEMF) on a Titan Krios microscope (Thermo Fisher) at 300 kV with a Gatan K3 detector in correlated double sampling super-resolution mode at a magnification of 105,000X corresponding to a pixel size of 0.415 Å. A total of 10,806 movies were collected; each movie was recorded for a total of 60 frames over 5.4 s with an exposure rate of 7.8 electrons/pixel/s. The datasets were collected using SerialEM software (14) with a defocus range of -0.9 and -2.4 µm.

A 24-hour data collection was performed on the best grid with Kap114/H2A-H2B/Nap1 FL/Ran$^{GTP}$ complex at the UTSW CEMF on a Titan Krios 300 kV microscope equipped with a Falcon 4 detector at a magnification of 165,000X at pixel size of 0.738 Å with defocus range of -0.9 to -2.2 μm. A total of 9331 movies were recorded for a total of 1114 frames over 3.6 s with an exposure rate of ~8 electrons/pixel/s.

**Cryo-EM data processing**

Cryo-EM data for the Kap114/H2A-H2B/Nap1 core complex was processed using cryoSPARC version 3 (15). The movies were subjected to Patch Motion Correction (binned twice) and Patch CTF Estimation. Training particles were obtained by blob picking on 100 micrographs and followed by 2D classification of the particle stack. Particles containing Kap114 were used to train on Topaz (16) which picked 665,443 particles. These particles were subjected to three rounds of 2D classification (300 classes/run). 113,471 particles that contained both Kap114 and Nap1 were split into two Ab-initio classes, the $Nap1_2$•H2A-H2B•Kap114 and $Nap1_2$•Kap114•H2A-H2B, and further classified using Heterogenous Refinement. 54,170 particles of $Nap1_2$•H2A-H2B•Kap114 and 59,310 particles of $Nap1_2$•Kap114•H2A-H2B were further refined using Non-Uniform (NU) Refinement resulting in 4.02 Å and 3.65 Å maps, respectively. To obtain more particles, these maps were used to generated templates for additional template picking on cryoSPARC. This resulted in 4,314,112 total particles that contained $Nap1_2$ only, Kap114•H2A-H2B, $Nap1_2$•H2A-H2B•Kap114 and $Nap1_2$•Kap114•H2A-H2B. After the first round of 2D classification ~1 million particles were further processed for Nap1 core dimer, and another ~1 million particles were further processed for the Kap114-containing complexes. Both Nap1 and Kap114 complexes underwent three more rounds of additional 2D classifications to obtain 468,627 and 675,417 particles, respectively. Two Ab-initio classes were generated for Nap1 particles, and four Ab-initio classes were generated for the Kap114 complexes followed by classification by Heterogeneous Refinement. 230,210 Nap1 particles were used in NU

Refinement resulting in the final map with 3.2 Å resolution. The $Nap1_2$•H2A-H2B•Kap114 and $Nap1_2$•Kap114•H2A-H2B particles obtained from Topaz were merged with ones obtained from template picking. Duplicates were then removed, resulting in 136,011 $Nap1_2$•H2A-H2B•Kap114 and 148,410 $Nap1_2$•Kap114•H2A-H2B final particles that were used for reconstruction in Non-uniform (NU) refinement with default parameters to obtain maps with 3.5 and 3.2 Å, respectively. Local refinement was performed using a custom fulcrum position at the center of h19 helices, determined in ChimeraX, to obtain improved maps for Nap1 in both reconstructions. Additionally, for $Nap1_2$•Kap114•H2A-H2B map, pose/shift gaussian prior during alignment was used.

Cryo-EM data for the Kap114/H2A-H2B/Nap1 FL/$Ran^{GTP}$ complex was processed using cryoSPARC version 4. The movies were subjected to Patch Motion Correction (no binning) and Patch CTF Estimation. Blob picking on all of the micrographs yielded 2,567,783 initial particles, which is followed by 5 rounds of 2D classification to 306,757 particles, which were submitted to Ab initio reconstruction and hetero-refinement to obtain 4 classes. Two classes containing Kap114 bound to $Ran^{GTP}$ were used as training particles for Topaz to pick 1,381,753 particles, which were cleaned by 3 rounds of 2D classification to 358,680 particles. These particles were submitted to Ab initio reconstruction and hetero-refinement to obtain 4 classes, one of which was used for further local CTF refinement and NU refinement with default parameters to obtain map with 2.9 Å refinement. Density for $Nap1_2$ and H2A-H2B was present in hetero-refinement map and early iterations for NU refinement, but not in the final map. Therefore, a mask was generated using a $Nap1_2$•H2A-H2B•Kap114 docked onto the low-resolution map and local refinement was performed with default parameters.

**Structure building/modeling**

The initial Nap1 model generated with AlphaFold-Multimer was used to build into the cryo-EM structure of the $Nap1_2$ (17). The initial models used to build the $Nap1_2$•Kap114•H2A-H2B structure included Kap114•H2A-H2B (PDB:8F0X), the Alpha-Fold model of Kap114 (AF-P53067-

F1), and our cryo-EM structure of the Nap1 dimer. The $Nap1_2$•H2A-H2B•Kap114 was built using the $Ran^{GTP}$•H2A-H2B•Kap114 cryo-EM structure (PDB: 8F1E), AF-P53067-F1, and our cryo-EM model of the Nap1 dimer as initial models. The $Nap1_2$•H2A-H2B•Kap114•$Ran^{GTP}$ structure was built using the $Nap1_2$•H2A-H2B•Kap114 cryo-EM structure determined in this study and the $Ran^{GTP}$•H2A-H2B•Kap114 (PDB: 8F1E) (10). All initial models were roughly docked into the map using UCSF Chimera or ChimeraX (18, 19) and then subjected to real-space refinement with global minimization and rigid body restraints in Phenix (20). The resulting structures were then manually rebuilt and refined using Coot (21), further corrected using ISOLDE (22) on UCSF ChimeraX and subjected to last round of refinement in Phenix. We used PDBe PISA to calculate solvent accessible surface areas (23). We also used PyMOL version 2.5 and the APBS electrostatic plugin for 3D structure and electrostatic analysis (24, 25).

**Fluorescence Polarization**

To generate fluorescently labelled Nap1 FL, the proteins were treated with 1 mM DTT for 30 min in room temperature and buffer exchanged into 20 mM HEPES pH 7.4, 500 mM NaCl, 10% (v/v) glycerol using a HiTrap Desalting column (Cytiva). 4 molar excess XFD488 (ATT Bioquest) was added and reaction incubated for 2 hrs at room temperature in the dark before removal of excess dye by SEC (Superdex S200 increase) in Assay Buffer. Fluorescence polarization assays were performed in Assay Buffer in triplicates of sixteen 20 µL samples. *Sc* H2A-H2B was serially diluted from 4 µM and mixed with 20 nM labelled $Nap1_2$ proteins in 1:1 ratio to yield final concentrations in a 384 well black bottom plate (Corning). Measurements were performed in a CLARIOstar Plus plate reader (BMG Labtech) with top optics using excitation filter 482-16, dichroic filter LP 504 and emission filter 530-40, 50 flashes per well. Gain was optimized to target mP of ~ 200. Data was analyzed in PALMIST (26) and plotted in GUSSI.

**SAXS methods**

To form the Nap1•Kap114 complex, a molar ratio of 1:3 $Nap1_2$ (core):Kap114 were mixed before purification using the Superdex200 Increase 10/300 column and the following buffer: 20 mM HEPES pH 7.4, 150 mM NaCl, 5% glycerol, and 1 mM TCEP. The protein complex, concentrated to 3 mg/mL, and the SEC buffer were both flash frozen to ensure precise buffer match. Measurements were taken with sample concentration between 1 – 3mg/mL at 10°C using the SAXS instrument at the 12-ID-B beamline of the Advanced Photon Source, Argonne National Laboratory. To eliminate all possible aggregates, samples were centrifuged at 15,000xg for 30 min, and only the top 50% of the solution was used for data measurement. To prevent potential radiation damage, samples were measured in a quartz capillary flow cell with a diameter of 1.5 mm under a constant flow rate of 10 µL/s. Each sample measurement was purged with its matching buffer solution. Data were collected using the Pilatus 2M area detector with a beam energy of 14k eV and beam current of 500 mA. 40 successive frames with an exposure of 1 s were recorded for each sample and the data was averaged for better signal/noise statistics. Scattering profiles were averaged, reduced, and merged from measurements at different concentrations using BioXTAS RAW software (27). Structural parameters and the distance distribution function, $P(r)$, were calculated with GNOM (28) and the ab initio electron density was reconstructed using DENSS (29).
